# Mechanistic Determinants of Slow Axonal Transport and Presynaptic Targeting of Clathrin Packets

**DOI:** 10.1101/2020.02.20.958140

**Authors:** Archan Ganguly, Florian Wernert, Sébastien Phan, Daniela Boassa, Utpal Das, Rohan Sharma, Ghislaine Caillol, Xuemei Han, John R. Yates, Mark H. Ellisman, Christophe Leterrier, Subhojit Roy

## Abstract

Clathrin has established roles in endocytosis, with clathrin-cages enclosing membrane infoldings, followed by rapid disassembly and reuse of monomers. However, in neurons, clathrin synthesized in cell-bodies is conveyed into axons and synapses via slow axonal transport; as shown by classic pulse-chase radiolabeling. What is the cargo-structure, and mechanisms underlying transport and presynaptic-targeting of clathrin? What is the precise organization at synapses? Combining live-imaging, mass-spectrometry (MS), Apex-labeled EM-tomography and super-resolution, we found that unlike dendrites where clathrin transiently assembles/disassembles as expected, axons contain stable ‘transport-packets’ that move intermittently with an anterograde bias; with actin/myosin-VI as putative tethers. Transport-packets are unrelated to endocytosis, and the overall kinetics generate a slow biased flow of axonal clathrin. Synapses have integer-numbers of clathrin-packets circumferentially abutting the synaptic-vesicle cluster, advocating a model where delivery of clathrin-packets by slow axonal transport generates a radial organization of clathrin at synapses. Our experiments reveal novel trafficking mechanisms, and an unexpected nanoscale organization of synaptic clathrin.

## INTRODUCTION

The cytosolic protein clathrin is an established player in receptor-mediated endocytosis. During this process, soluble clathrin is recruited to the plasma membrane, forming clathrin-coated pits and coated vesicles. After internalization of coated vesicles, the clathrin coat is rapidly removed by uncoating proteins like auxilin and heat shock protein 70 (Hsc70), and the released clathrin-monomers are reused for subsequent rounds of endocytosis. This ‘clathrin-coated vesicle cycle’ has been studied for over forty years, and a host of adapters and regulators are known to be involved. Clathrin also plays roles in sorting of intracellular membranes, although this is less studied (Jung and Haucke, 2007; McMahon and Boucrot, 2011; Traub and Bonifacino, 2013).

In neurons, clathrin synthesized in cell-bodies is transported into axons, enriching at presynapses. Previous pulse-chase radiolabeling studies in vivo have established that clathrin is conveyed in slow axonal transport (Black et al., 1991; de Waegh and Brady, 1989; Elluru et al., 1995; Garner and Lasek, 1981; Gower and Tytell, 1987). Slow transport is a mysterious rate-class carrying cytosolic (or ‘soluble’) proteins to the axon-tips and presynaptic terminals over days to months in vivo, unlike fast vesicular cargoes that are rapidly transported in minutes to hours; reviewed in (Maday et al., 2014; Roy, 2014). Although radiolabeling studies have established that clathrin is conveyed in slow axonal transport, the methods used could only infer overall movement by determining the changing radioactivity-pattern in axons over time, and many questions remain. How can clathrin undergo organized slow axonal transport, given the ephemeral nature of cytoplasmic clathrin-assemblies? Indeed, previous studies in cultured hippocampal neurons showed rapid on/off behavior of GFP-tagged clathrin puncta in dendrites, lasting only for seconds (Blanpied et al., 2002). What does the axonal cargo-structure of clathrin look like? What are the underlying mechanisms conveying clathrin in slow transport? How is clathrin targeted to presynapses and retained there? What is the nanoscale organization of clathrin at synapses? The precise function of clathrin at presynapses is also unclear (Chanaday et al., 2019; Chanaday and Kavalali, 2018; Milosevic, 2018; Royle and Lagnado, 2010). Although cytosolic slow axonal transport remains enigmatic, recent studies from us and others using cultured neurons as a model-system – to visualize and manipulate slow transport – have begun to offer some answers (Chakrabarty et al., 2019; Ganguly et al., 2017; Scott et al., 2011; Tang et al., 2013; Twelvetrees et al., 2016). Here we explored mechanisms underlying clathrin transport using a multifaceted approach involving live imaging, super-resolution, 3D EM and proteomics. Our experiments reveal previously unknown long-lasting axonal clathrin assemblies in axons that are unrelated to endocytosis and seem specialized for cargo-delivery to synapses. Collectively, the data uncover mechanistic details underlying the slow axonal transport and presynaptic targeting of clathrin – that has been unclear for decades – and reveal a novel nanoscale organization in axons and synapses.

## RESULTS

### Dynamic clathrin transport-packets mediate slow axonal transport

**Figure 1A** shows the distribution of endogenous clathrin in cultured hippocampal neurons. Though the overall appearance of clathrin-puncta seemed similar in somatodendritic and axonal compartments by immunostaining, their dynamics were different. To visualize clathrin in living neurons, we used GFP (or mCherry) tagged to the clathrin light chain (CLC) – a structurally and functionally active fusion protein (Gaidarov et al., 1999) that has been used in many other studies (Blanpied et al., 2002; Pelassa et al., 2014; Zhao and Keen, 2008). As reported previously (Blanpied et al., 2002), GFP:CLC showed largely on/off dynamics in dendrites, likely reflecting transient clathrin recruitment to coated-pits/vesicles (kymograph in **Fig. 1B,** also see **Supp. Movie 1**). Interestingly however, axonal GFP:CLC particles were more long-lasting, and a significant fraction (~ 40%) were motile; moving with an overall bias towards the axon-tip (representative kymograph and quantification in **Fig. 1C-E**, also see **Supp. Movie 2**). Two-color live imaging of GFP:CLC and synaptophysin:mRFP revealed clear instances where clathrin particles were transported independently of synaptophysin (**Supp. Fig. 1**). To distinguish the moving axonal clathrin structures from dendritic clathrin-coated pits and other coated vesicles that may be involved in endocytosis, here we refer to them as ‘clathrin transport-packets’.

**Figure 1:**
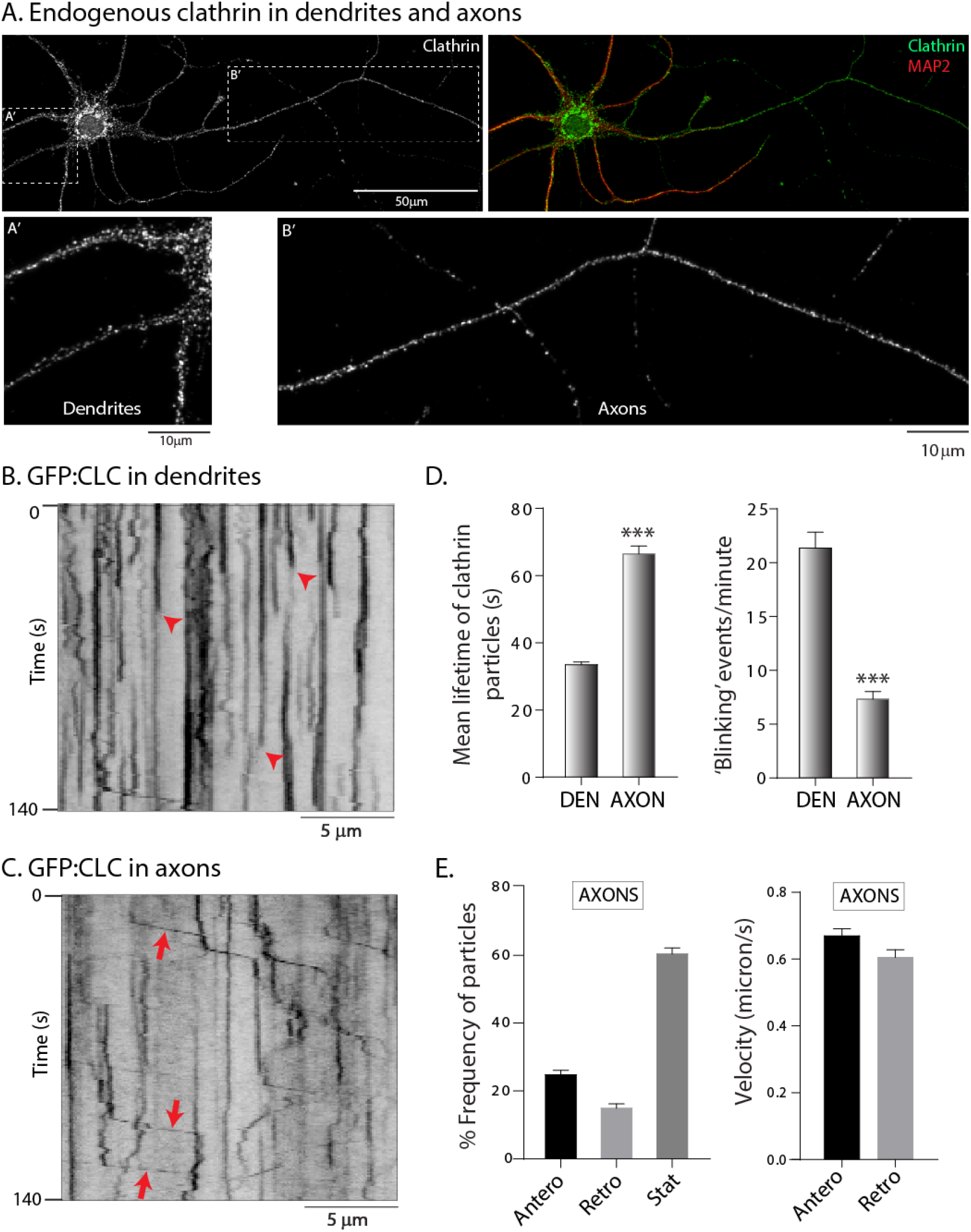
Differential dynamics of clathrin in dendrites and axons. (A) Clathrin and MAP2 immunostaining in cultured hippocampal neurons. Note punctate clathrin distribution in both somatodendritic and axonal compartments. (B) Kymograph of GFP:CLC dynamics in a dendrite. Note abrupt appearance and disappearance of fluorescence (some marked by red arrowheads), indicating assembly/disassembly of clathrin in coated-pits (receptor mediated endocytosis). (C) Kymograph of GFP:CLC dynamics in an axon. Note rapid, infrequent, and anterogradely-biased movement of clathrin particles (some marked by red arrows). (D) Mean lifetimes of GFP:CLC particles – as determined from kymographs – were significantly higher in axons, while ‘blinking events’ – representing clathrin assembly/disassembly and receptor mediated endocytosis is much lower in axons, compared to dendrites (1113 particles from 22 dendrites and 403 particles from 41 axons were analyzed; data was pooled from 3 independent experiments; ***p<0.0001) (E) About 40% of the axonal GFP:CLC particles were mobile, with a larger fraction of particles moving anterogradely. Note that the velocity of particles is consistent with motor-driven cargoes (460 anterograde and 263 retrograde events were analyzed from 41 axons; data was combined from 3 independent experiments).

Previously, we developed an assay to visualize the anterogradely-biased transport of cytosolic cargoes moving in slow transport – called the “intensity-center shift” assay (Roy et al., 2012; Scott et al., 2011). Here, cultured neurons are transfected with photoactivatable GFP (PAGFP) tagged to the cytosolic protein of interest, and a discrete axonal population of tagged cytosolic-proteins – typically within ~ 30μm of axon-length – is photoactivated with 405nm light (schematic in **Fig. 2A**). Thereafter, a line is drawn along the longitudinal axis of the photoactivated zone, and the intensity-center (or centroid) of fluorescence is monitored over time. While untagged PAGFP rapidly diffuses without any intensity-center shift, cytosolic proteins disperse with a slow anterogradely-biased shift; consistent with slow transport (Chakrabarty et al., 2019; Ganguly et al., 2017; Scott et al., 2011; Tang et al., 2012; Tang et al., 2013; Twelvetrees et al., 2016). An exemplary kymograph from these experiments is shown in **Figure 2B**, along with the ensuing raw and ensemble data from these experiments (**Fig. 2C-D**). Note that PAGFP:CLC, but not untagged PAGFP, has an overall anterograde bias with a predicted average rate of ~ 0.006 μm/s or ~ 0.5 cm/day, consistent with the overall rates of slow transport in radiolabeling studies (Garner and Lasek, 1981). To further confirm that the biased transit was due to kinetics of axonal clathrin particles – and also to verify the results using a complementary assay – we first visualized green Dendra2:GFP:CLC puncta in axons and then photoconverted them to red (**Fig. 2E**, top). Intensity-center analysis of the red fluorescence also showed an anterograde bias (**Fig. 2E**, bottom). Collectively, the data indicate that the intermittent movement of axonal clathrin transport-packets generates an overall anterograde bias of the clathrin population, at rates consistent with slow axonal transport.

**Figure 2:**
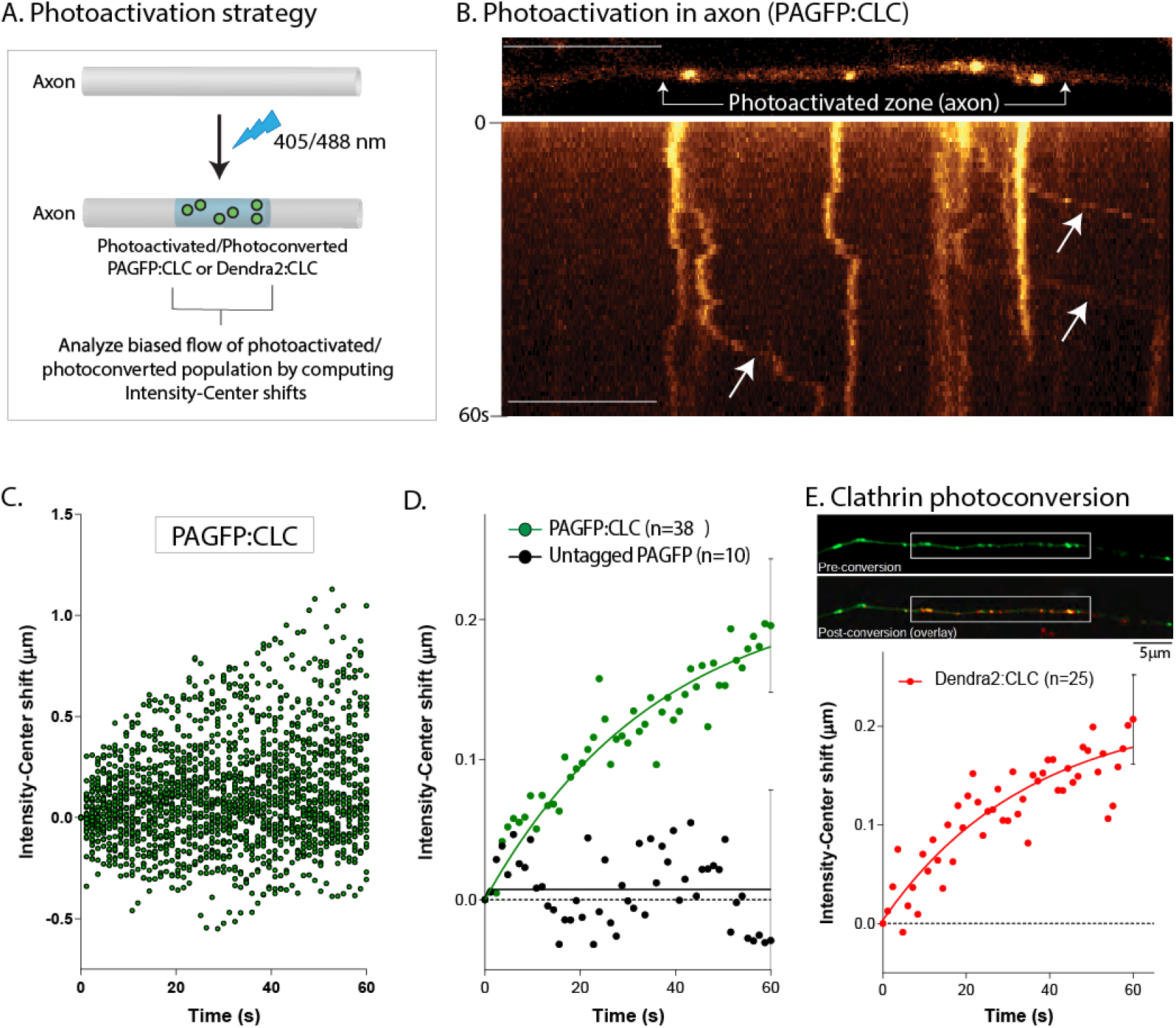
Slow anterograde bias of clathrin in axons. (A) Strategy for analyzing biased axonal transport of clathrin in axons. Clathrin is photoactivated (or photoconverted) in a discrete portion (~30 μm) of the axon, and dispersion of the fluorescent clathrin population is tracked over time. An anterograde shift in the center of fluorescence (intensity-center shift) reflects the biased movement of the clathrin population in the axon (see results and methods for more details). (B) Kymograph from a photoactivation (PAGFP:CLC) experiment. Note anterograde movement of photoactivated clathrin particles (arrows). Scale bars=10 microns. (C) Raw data of intensity-center shifts from all PAGFP:CLC experiments in axons. Note anterograde drift of datapoints (38 neurons from 3 separate cultures were analyzed). (D) Ensemble data showing mean intensity-center shift of PAGFP:CLC and PAGFP-only in axons, with fitted curves. Note that while there is an overall anterograde bias of PAGFP:CLC, there is no bias with untagged PAGFP-only (10 neurons from 2 separate cultures were analyzed for PAGFP-only). (E) Neurons were transfected with Dendra2:CLC, axonal clathrin particles were identified by green fluorescence, and intensity-center shifts of photoconverted (red) particles was analyzed. Note anterograde shift of the particles, indicating that the movement of clathrin particles generate the bias (25 neurons from 3 separate cultures were analyzed).

### Dynamics of axonal clathrin transport-packets is independent of endocytosis

The persistence and biased kinetics of the transport-packets suggest that these may represent a neuronal organelle specialized for axonal transport of clathrin, with no role in endocytosis. To test this idea, we asked if either global inhibition of endocytosis, or specific attenuation of clathrin-dependent endocytosis, would also affect the motility of axonal transport-packets. First, we globally suppressed endocytosis in cultured neurons using Dynasore – a small-molecule inhibitor of dynamin (Das et al., 2013; Macia et al., 2006). Next, we also specifically inhibited clathrin-dependent endocytosis with CHC-T7-Hub, a clathrin heavy-chain mutant known to disrupt clathrin-dependent endocytosis in a dominant-negative fashion (Bennett et al., 2001). As reported previously in non-neuronal cells, the CHC-T7-Hub mutant attenuated receptor-mediated endocytosis in cultured hippocampal neurons (**Supp. Fig. 2A-B**). As expected, the on/off dynamics of GFP:CLC in dendrites was essentially abolished in neurons treated with Dynasore (**Fig. 3A)**, or transfected with the CHC-T7-Hub mutant (**Supp. Fig. 2C-D**). Interestingly however, neither Dynasore, nor the CHC-T7-Hub mutant had any effect on the mobility of axonal clathrin transport-packets (**Fig. 3B-D**). Since the motility of individual clathrin transport-packets generate the overall anterogradely-biased flow of the axonal clathrin population (see **Fig. 2E**), a prediction is that endocytosis-inhibition would not interfere with the slow transport of clathrin. Indeed, neither Dynasore nor the CHC-T7-Hub mutant had any effect on the biased transit of PAGFP:CLC in axons (**Fig. 3E**). Previous studies suggest that the slow transport of is microtubule-dependent (Scott et al., 2011; Twelvetrees et al., 2016). Low-levels of the microtubule-disrupting drug Nocodazole – that completely blocks vesicle transport see **Supp. Fig. 2E** – also inhibited movement of clathrin transport-packets (**Fig. 3F**). Taken together, the data indicate that axonal clathrin transport-packets are unique conveyance structures that are unrelated to endocytosis.

**Figure 3:**
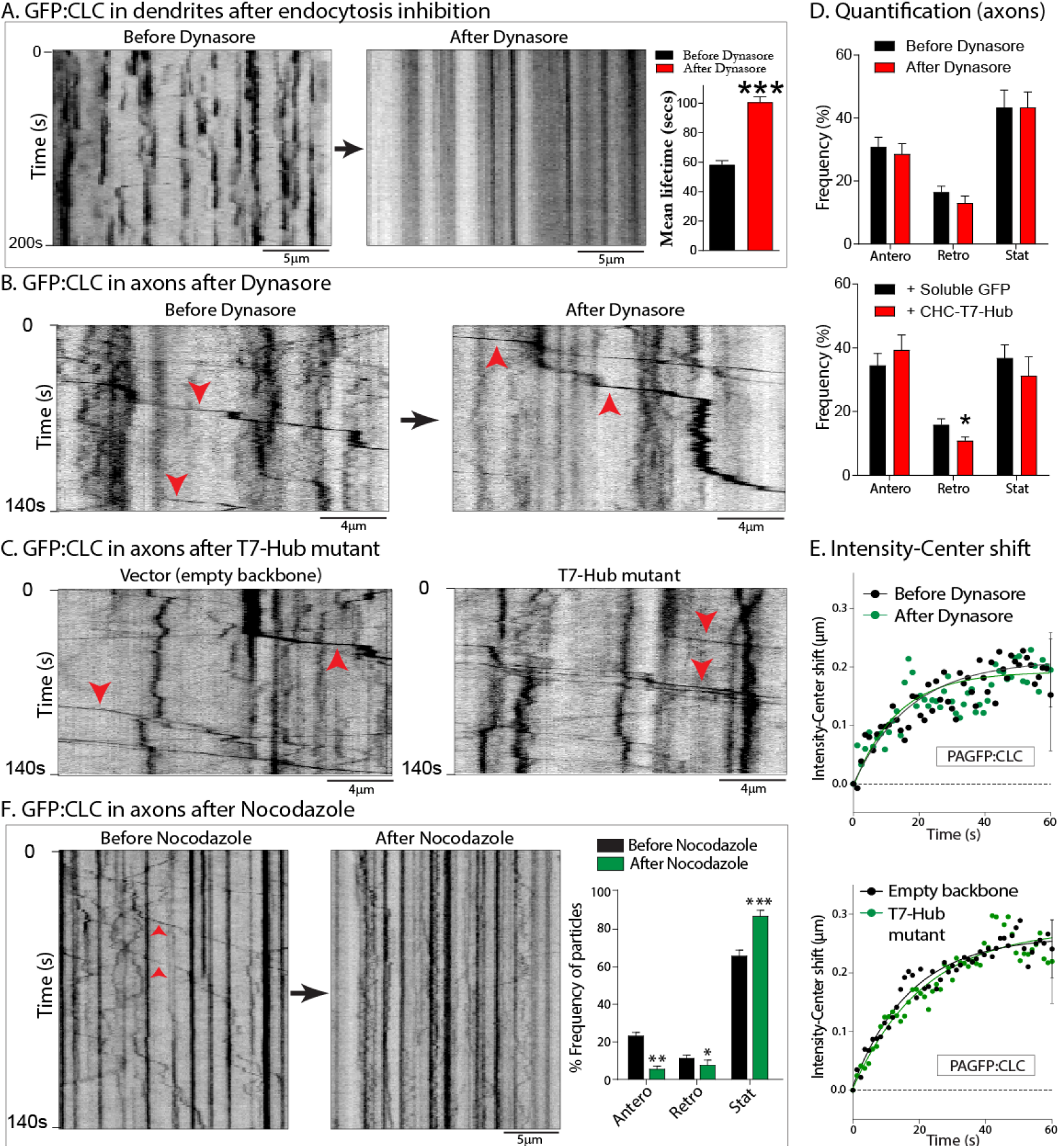
Motility of axonal clathrin transport-packets is independent of endocytosis, but microtubule-dependent. (A) Kymographs of GFP:CLC dynamics from the same dendrite before and after inhibition of endocytosis (Dynasore). Note that the blinking events – reflecting endocytosis – are abolished after drug-treatment. GFP:CLC fluorescence-lifetimes are quantified on right (~ 150-250 particles from 10 neurons were analyzed; data from three separate cultures; p<0.0001) (B) Kymographs of GFP:CLC dynamics from the same axon before and after Dynasore treatment. Note that vectorial movement of clathrin transport-packets (some marked by arrowheads) continue after drug-treatment. (C) Kymographs of axonal mCherry:CLC dynamics from neurons transfected with a dominant-negative clathrin mutant (T7-Hub) that specifically interferes with clathrin-dependent endocytosis (see **Supp. Fig. 2**); or control. Note that vectorial movement of clathrin transport-packets (some marked by arrowheads) are similar in the two groups. (D) Quantification of all mCherry:CLC axonal transport data – with and without endocytosis inhibition – global or clathrin-dependent. (~ 140-170 motile particles were analyzed from 10-12 neurons; data from 2-3 separate cultures; *p<0.01) (E) Ensemble intensity-center shift data (PAGFP:CLC slow transport assay) from neurons treated with Dynasore or co-transfected with the T7-Hub mutant. Note that inhibiting global – or clathrin-dependent – endocytosis has no effect on the slow anterograde bias of the clathrin population in axons (20-30 axons from two and four separate cultures were analyzed). (F) Kymographs of GFP:CLC dynamics from the same axon before and after treatment with 10μg/ml Nocodazole for 30 min., with quantification on right. Note that motility of the clathrin transport-packets is attenuated after drug-treatment.

### Structure of clathrin transport packets

What do the transport-packets look like? To visualize the ultrastructure of axonal clathrin, we tagged clathrin to Apex – an engineered peroxidase that acts as an EM-tag by catalyzing the polymerization and local deposition of diaminobenzidine (DAB); subsequently recruiting electron-dense osmium for EM contrast (Martell et al., 2012). Cultured neurons were transfected with APEX:GFP:CLC, fixed and incubated with DAB and H_2_O_2_ (see schematic in **Fig. 4A** and methods). Transfected neurons were visualized by GFP fluorescence and DAB-staining; dendrites and axons were identified by morphology; and the same neurons and processes were visualized by EM tomography (see **Supp. Fig. 3** and methods). As expected, coated pits were readily seen in soma and dendrites (**Supp. Fig. 3B**). However, axons contained intact clathrin-coated structures that were inside the axon-shaft, with no obvious association with the axonal plasma membrane, and were also frequently adjacent to microtubules, as seen by EM-tomography (**Fig. 4B-B”**; also see 3-D view in **Supp. Movies 3 and 4**). Such intact clathrin-structures are surprising, given the transient nature of clathrin, and we reason that these represent transport-packets. The diameter of these clathrin-structures as determined by EM was ~ 50nm and ~125nm (spherical core and outer diameter respectively); consistent with previous cryo-EM studies of in vitro reconstituted clathrin (Heymann et al., 2013).

**Figure 4:**
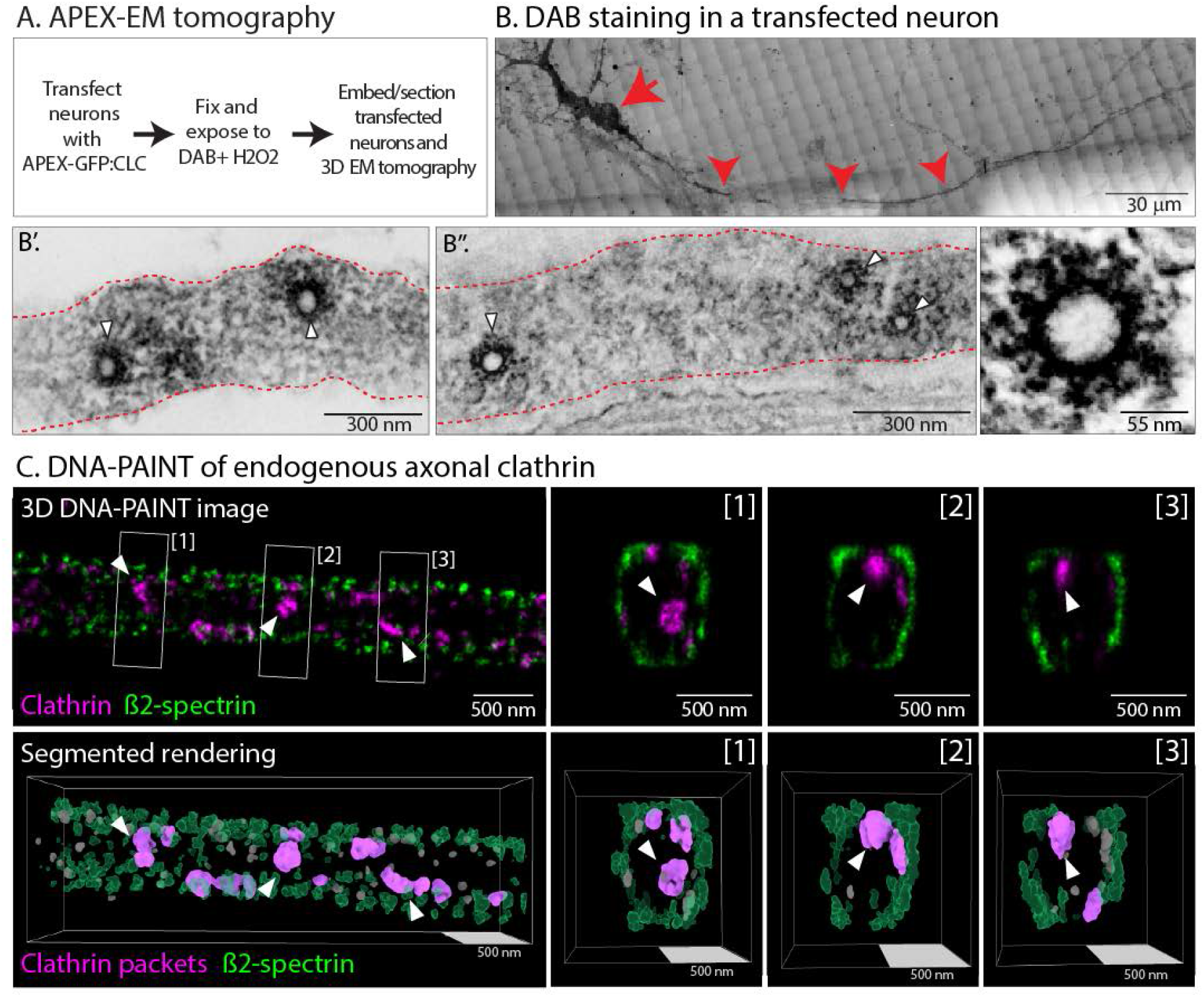
Ultrastructure and super-resolution-imaging of axonal clathrin transport-packets. (A) Strategy to visualize clathrin by Apex-labeling and EM-tomography. Neurons were transfected with Apex:GFP:CLC – an EM-tag inducing local deposition of DAB, leading to dense contrast (also see results and methods). (B) A DAB-stained neuron identified on gridded membrane (soma and axon marked by arrow and arrowhead respectively; also see methods and **Supp. Fig. 3**). (B’ and B”) EM of the axon above (a single plane of tomogram shown). Note unequivocal clathrin-coated organelles in the axon-shaft (white arrowheads), with average diameters of ~ 50 nm and ~125 nm for the spherical core and outer diameter respectively. (C) DNA-PAINT of endogenous clathrin in the axon shaft. **<u>Top panels</u>**: β2-spectrin (green, to mark axonal boundaries) and clathrin heavy chain (magenta). An X-Y image is shown on the left, along with three Z-slices on right ([1]-[3]). **<u>Bottom panels</u>**: Segmented rendering from above 3D-PAINT images (small excluded particles are in grey). Equivalent sphere diameters for the clathrin particles marked by arrowheads: 79, 70 and 70 nm for particles highlighted in [1], [2] and [3] respectively.

We next used DNA-PAINT, a single-molecule based super-resolution microscopy method (Jungmann et al., 2014) to visualize the nanoscale organization of endogenous clathrin along axons. Axonal boundaries were determined by visualizing β2-spectrin, a component of the submembrane periodic scaffold along axons [reviewed in (Leterrier et al., 2017)]. Unlike the tedious Apex EM-imaging process involving light/EM correlation of transfected axons, DNA-PAINT allowed us to visualize and characterize a large number (>500) of axonal clathrin structures. From 3D multicolor DNA-PAINT images of clathrin and ß2-spectrin, we generated 3D-renderings for the segmentation and measurement of axonal clathrin particles; some of which are likely to be transport-packets (**Fig. 4C**). The average equivalent diameter of clathrin heavy-chain labeled packets was found to be 47 nm, consistent with the core size of the structures seen in our EM data. Interestingly, most clathrin packets seen by super-resolution-imaging were apposed to β2-spectrin, suggesting association with the submembrane actin/spectrin scaffold (see cross-sections of axons in **Fig. 4C** and **Supp. Movie 5)**. Given that the majority of axonal clathrin particles are immobile at any given time, this may reflect a possible anchoring of clathrin assemblies to the periodic actin/spectrin lattice (also see data with actin/myosin-6 later).

### Composition of axonal clathrin and transport mechanisms

To define the composition of axonal clathrin, we performed immunoprecipitation (IP) of clathrin from mouse sciatic axon fractions, followed by MUDPIT-MS [multidimensional protein identification technology mass spectrometry; see (Liao et al., 2008; Yates et al., 2009)], as shown in **Figure 5A**. As expected, clathrin was one of the most abundant peptides isolated (**Fig. 5B-C**). A diverse group of cytosolic proteins were identified by MS, belonging to several functional groups (**Fig. 5D**; see **Supp. Table 1** for full list). Network analyses revealed two interesting features. First, the axonal clathrin-interactome consisted of proteins known to disassemble clathrin coated-pits (such as Hsp70, GAK, synaptojanin), as well as proteins related to pit-formation (such as dynamin, AP-complex). Second, a large group of cytoskeleton-associated proteins – including actin and myosins – were seen in the datasets (**Fig. 5D**). Previous studies have suggested that axonal actin accumulations may act as a mesh for vesicles moving in fast transport (Sood et al., 2018), and an unconventional minus-end directed myosin – myosin-6 – has also been implicated as an anchor for some axonal cargoes (Lewis et al., 2011). Moreover, actin and clathrin have well-established anatomic and functional links in non-neuronal cells [reviewed in (Kaksonen et al., 2006)].

**Figure 5:**
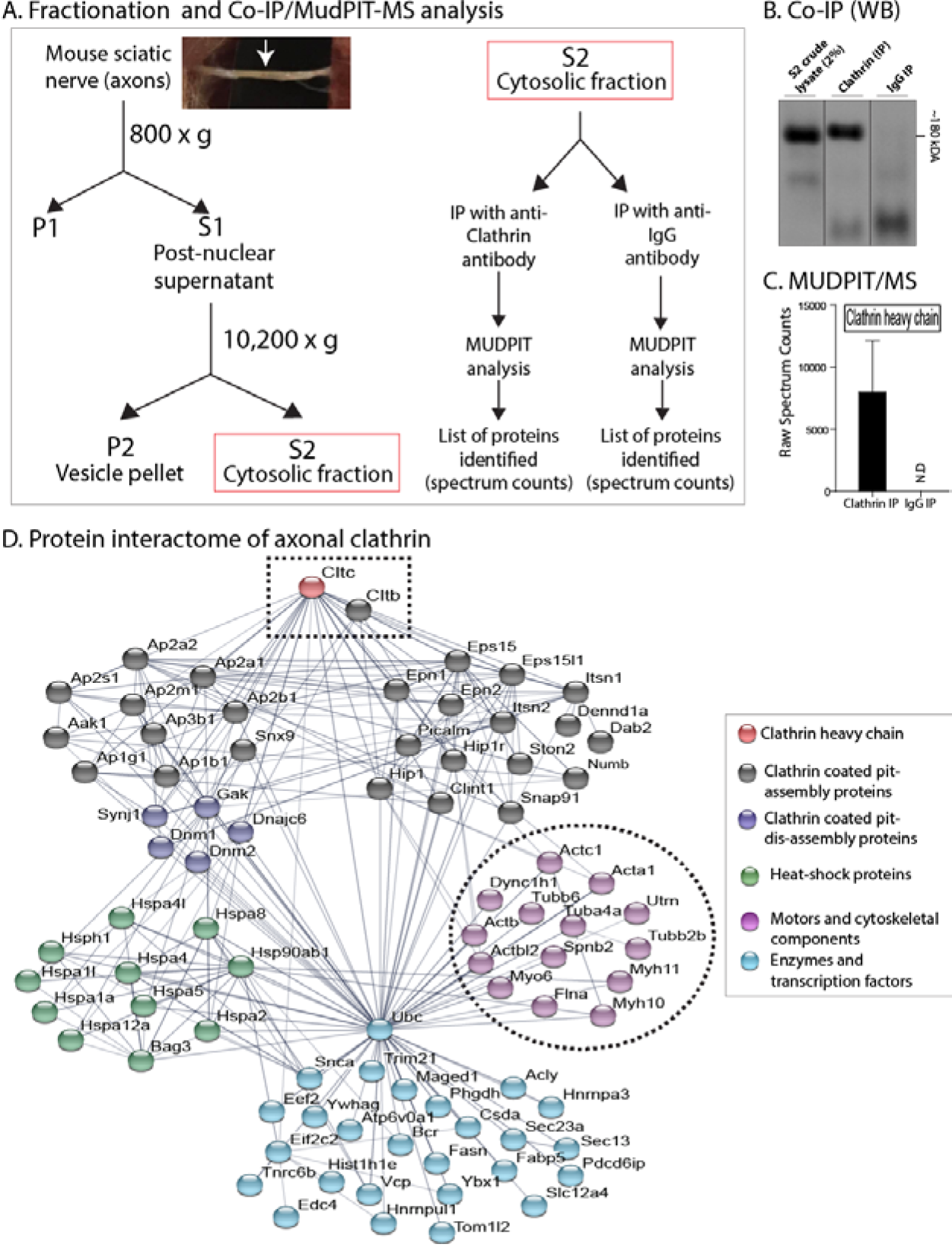
The axonal clathrin interactome. (A) Schematic for fractionation and Mudpit-MS. Clathrin was immunoprecipitated from cytosolic (S2) sciatic nerve fractions (dissected from mice, see inset picture and arrow), and associating proteins were identified by MS. (B, C) Western blots showing clathrin in IP fractions, also confirmed by high spectrum counts in MS-data. (D) An interactome of clathrin-associated proteins (clathrin marked with rectangle), grouped into functional clusters. The ‘actin-hub’ (circled) was of particular interest as previous studies have established mechanistic links between clathrin and actin (also see results).

Thus, we wondered if actin and myosin-6 might play a role in anchoring clathrin transport-packets in axons. Towards this, we first visualized axonal clathrin and actin using two-color live imaging. In previous studies, using a probe for filamentous actin (GFP tagged to the calponin-homology domain of utrophin, GFP:Utr-CH), we found that axon-shafts have focal “hotspots” every ~ 3-4μm along its length, where actin assembles and disassembles continuously, and that these foci serve as nidus for nucleation of actin filaments that elongate bidirectionally along the axon-shaft [“actin trails”, see (Ganguly et al., 2015)]. Interestingly, there was a striking colocalization of actin hotspots and clathrin in axons, though the actin foci were much more short-lived, as expected (**Fig. 6A-A’**; more examples in **Supp. Fig. 4**). Also, disruption of myosin-6 using a dominant-negative mutant (Lewis et al., 2011) led to a marked increase in the mobility of the GFP:CLC labeled transport-packets (**Fig. 6B**; quantified in **Fig. 6C**). Interestingly, myosin-6 disruption did not affect either the fast transport of synaptophysin-carrying vesicles, or the axonal dynamics of synapsin (**Supp Fig. 5A-D**) – another slow-transport cargo (Scott et al., 2011; Tang et al., 2013). Correspondingly, pharmacologic or genetic inhibition of Hsc70 – known to block the biased transit of synapsin in axons of cultured neurons (Ganguly et al., 2017) – had no effect on the axonal GFP:CLC motility (**Supp Fig. 5E**). Thus, the data imply that the mechanistic basis for the transport of two slow-component cargoes – clathrin and synapsin – are different. Finally, disruption of either actin or myosin-6 also led to an increase in the slow anterograde-bias of the PAGFP:CLC population in axons, as determined by the intensity-center shift assay (**Fig. 6D**). Collectively, the data suggest that actin and myosin-6 play a cooperative role in anchoring clathrin transport-packets along the axon – possibly contributing to its sluggish movement in slow transport.

**Figure 6:**
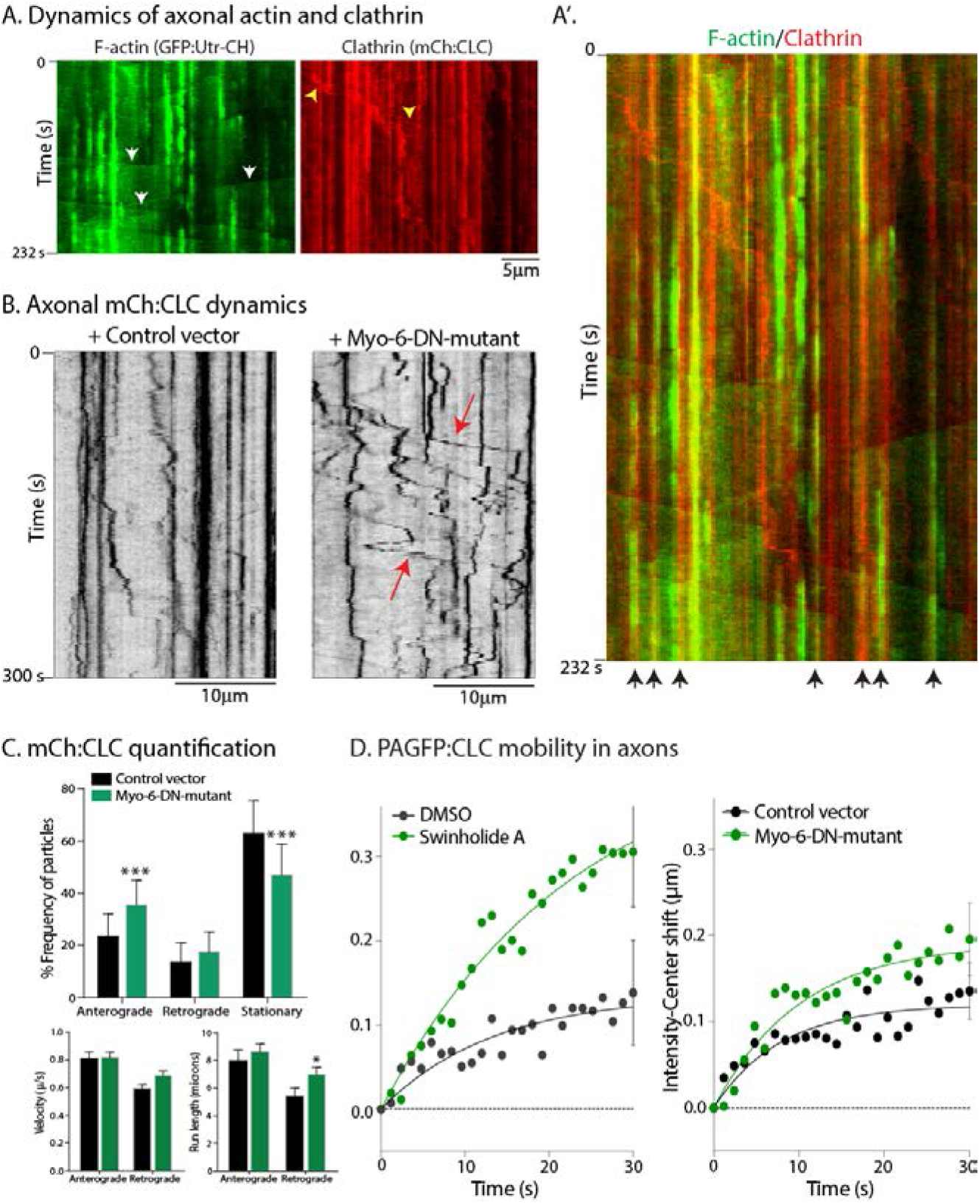
Actin and Myosin-6 as putative tethers of clathrin transport. **(A)** Neurons were transfected with GFP:Utr-CH – a marker of filamentous actin (F-actin) – and mCh:CLC; and both fluorophores were visualized by near-simultaneous live imaging (see methods). Kymographs show actin ‘hotspots’ and ‘trails’ (latter marked by white arrowheads) as described previously (see results). Some clathrin transport-packets are marked by yellow arrowheads. Note colocalization of F-actin and clathrin in the green/red overlaid kymograph (A’, marked by black arrows, bottom). **(B)** Kymographs of axonal GFP:CLC dynamics from neurons transfected with a dominant-negative myosin-6 inhibitor (Myo-6-DN) or vector-controls. Note enhanced motility of clathrin transport-packets after myosin-6 disruption; quantified in **(C)**. (~ 200-300 mobile particles from 20-24 axons were analyzed; data from four separate cultures. (***p<0.0001). **(D)** Mean intensity-center shifts of PAGFP:CLC in axons (slow transport assay) treated with the actin-disrupting agent Swinholide (left); or in neurons transfected with Myo-6-DN mutant. Note that the slow, anterogradely-biased flow of clathrin is upregulated upon disruption of actin or myosin-6 (15-25 axons were analyzed from three separate cultures).

### Delivery and organization of presynaptic clathrin packets

Previous studies have demonstrated enrichment of clathrin at presynaptic boutons and terminals, though its precise function at this locale has been controversial in recent years (Chanaday et al., 2019; Milosevic, 2018; Watanabe et al., 2013). While our live imaging of GFP:CLC in cultured neurons showed clathrin accumulation at boutons, we noticed that the localization was not homogenous – like classical synaptic markers – but appeared as multiple motile (‘jiggling’) puncta within single boutons (**Fig. 7A-A’**; also see **Supp. Movie 6**). Also, as seen in the kymograph (**Fig. 7A”**), we saw motile clathrin puncta targeting to boutons. Due to the constant mobility of the synaptic clathrin particles within micron-sized boutons – and subsequent overlapping of fluorescent puncta – we could not confidently track each particle long enough, to determine their lifetimes. However our impression was that the GFP:CLC particles did not behave like the ‘blinking dots’ at dendrites, but were more long-lasting like the axonal clathrin-particles (see kymograph in **Fig. 7A”**). The morphology and dynamics of synaptic clathrin puncta suggest that axonal transport and subsequent presynaptic “trapping” of a discrete number of transport-packets is the basis of clathrin enrichment at boutons. We thus asked if the total fluorescence of clathrin at individual boutons was an aggregate of integer-multiples of axonal puncta/transport-packets. Overlaid histogram-distributions of GFP:CLC fluorescence intensities in axons (green) and presynapses (red) are shown in **Figure 7B** – top panel. Note that while the intensities of axonal (non-synaptic) GFP:CLC occupy a single-peak, synaptic GFP:CLC intensities are higher, and distributed as multiple peaks. **Figure 7B** – middle panel shows predicted fluorescence peaks of synaptic clathrin as integer-multiples of the axonal GFP:CLC fluorescence intensity peak (which is assumed as ‘x1’). From this quantification of presynaptic clathrin fluorescence (**Fig. 7B** – bottom), single boutons are predicted to contain ~ 4-8+ clathrin-packets.

**Figure 7:**
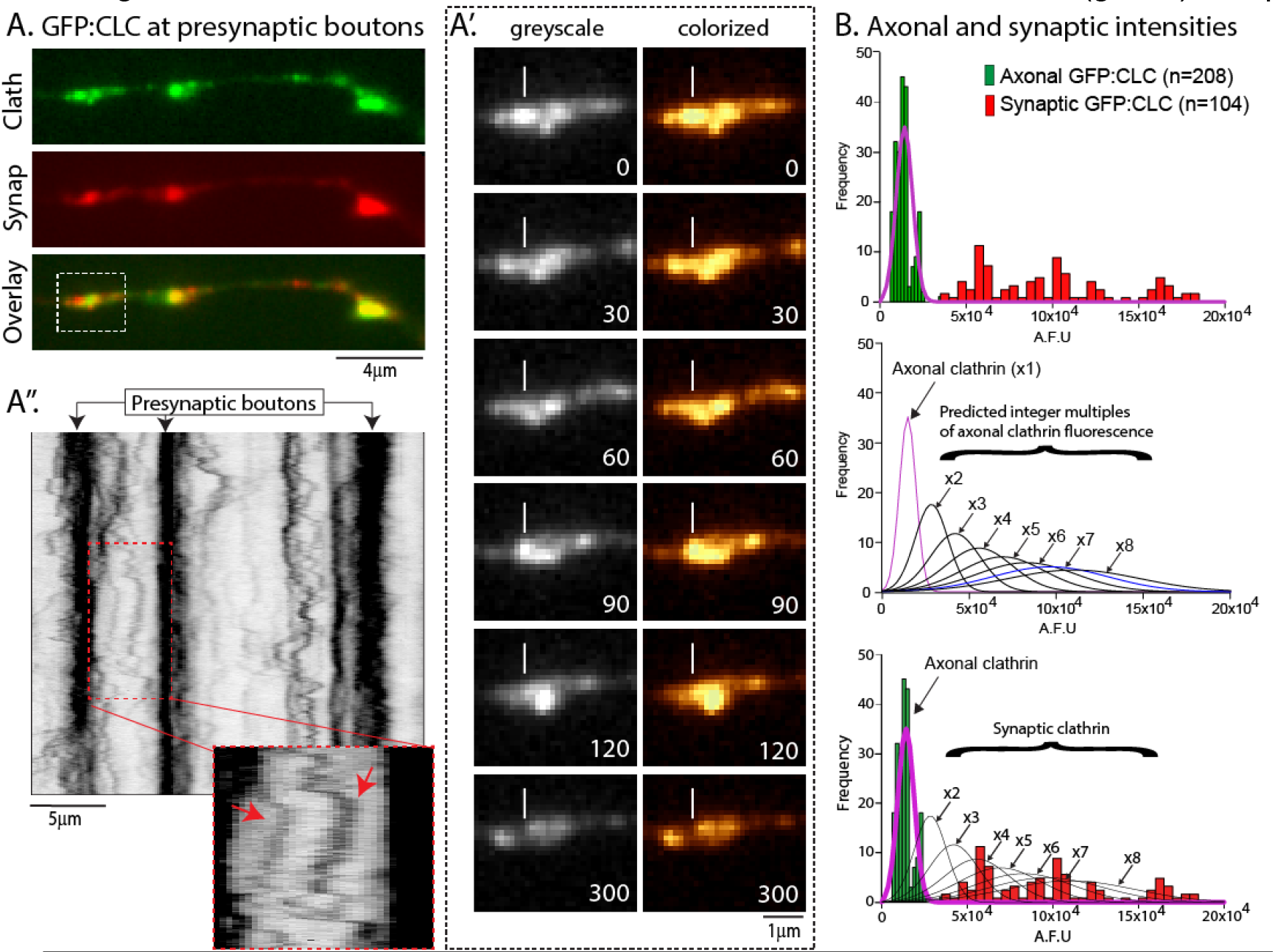
Synaptic targeting of integer-numbers of axonal clathrin transport packets. (A) Neurons were co-transfected with GFP:CLC (above) and synaptophysin:mCh (middle); distal boutons were located using synaptophysin (see methods). Panels show images of clathrin and synaptophysin at boutons. (A’) GFP:CLC dynamics in bouton from boxed region in (A). Note multiple motile clathrin puncta within individual boutons (small vertical line marks a reference-point in image; elapsed time in seconds on lower right). (A’’) Kymograph of (A). Note multiple GFP:CLC particles moving between – and targeting to – boutons (arrows; also see results). (B) **<u>Top</u>**: Histogram showing distribution of GFP:CLC intensities in axons and synapses. Note that while axonal clathrin intensities are tightly clustered in a single peak (green), synaptic intensities are distributed as multiple peaks of fluorescence (red). **<u>Middle</u>**: Mean fluorescence of axonal clathrin was considered as 1x, and hypothetical integer-multiple curves were generated (fitted curves shown). **<u>Bottom</u>**: Axonal and synaptic GFP:CLC fluorescence data were overlaid with the curves shown in middle panel. Note that data predicts that the synaptic clathrin fluorescence is composed of 4-8+ integer-multiples of axonal fluorescence (100-200 particles were analyzed from three separate cultures).

The particle-like presynaptic appearance of clathrin is somewhat surprising, considering its established role in mediating endocytosis at the plasma membrane. To clarify the spatial relationship of endogenous presynaptic clathrin to synaptic-vesicle clusters, we visualized synapses and adjacent axons at nanometer resolution in 3-D, with two-color DNA-PAINT; using antibodies to clathrin and synapsin, known to bind synaptic-vesicles (see methods for details). **Figure 8A-A”** shows an axon from these experiments. Note that multiple clathrin particles (magenta) are seen abutting the synaptic-vesicle cluster (green). Further 3-D views (**Fig. 8B**) unequivocally show that the clathrin particles are circumferentially organized around the synaptic-vesicle cluster, and rarely seen deep within the cluster (also see **Supp. Movie 7**). Analysis of the segmented rendering data show that each presynapse contains ~ 4-12 clathrin particles (**Fig. 8C**) that were on average ~ 43 nm in equivalent diameter (**Fig. 8D**), consistent with the size of clathrin assemblies seen in the axon-shaft (see **Fig. 4**). The morphologic similarities between the axonal and presynaptic clathrin particles suggest that they are related to transport-packets. The large number of clathrin particles visualized by DNA-PAINT (>1200 packets from >100 synapses) also allowed us to explore correlations between clathrin-packets and synaptic-vesicle clusters. Interestingly, the total volume of all clathrin-packets at a given synapse correlated with the total volume of the synaptic-vesicle cluster (**Fig. 8E**), suggesting functional links between clathrin-levels and synaptic-vesicle organization. Taken together, the data suggest that motile axonal transport-packets are targeted to presynapses, where they accumulate around the synaptic vesicle cluster.

**Figure 8:**
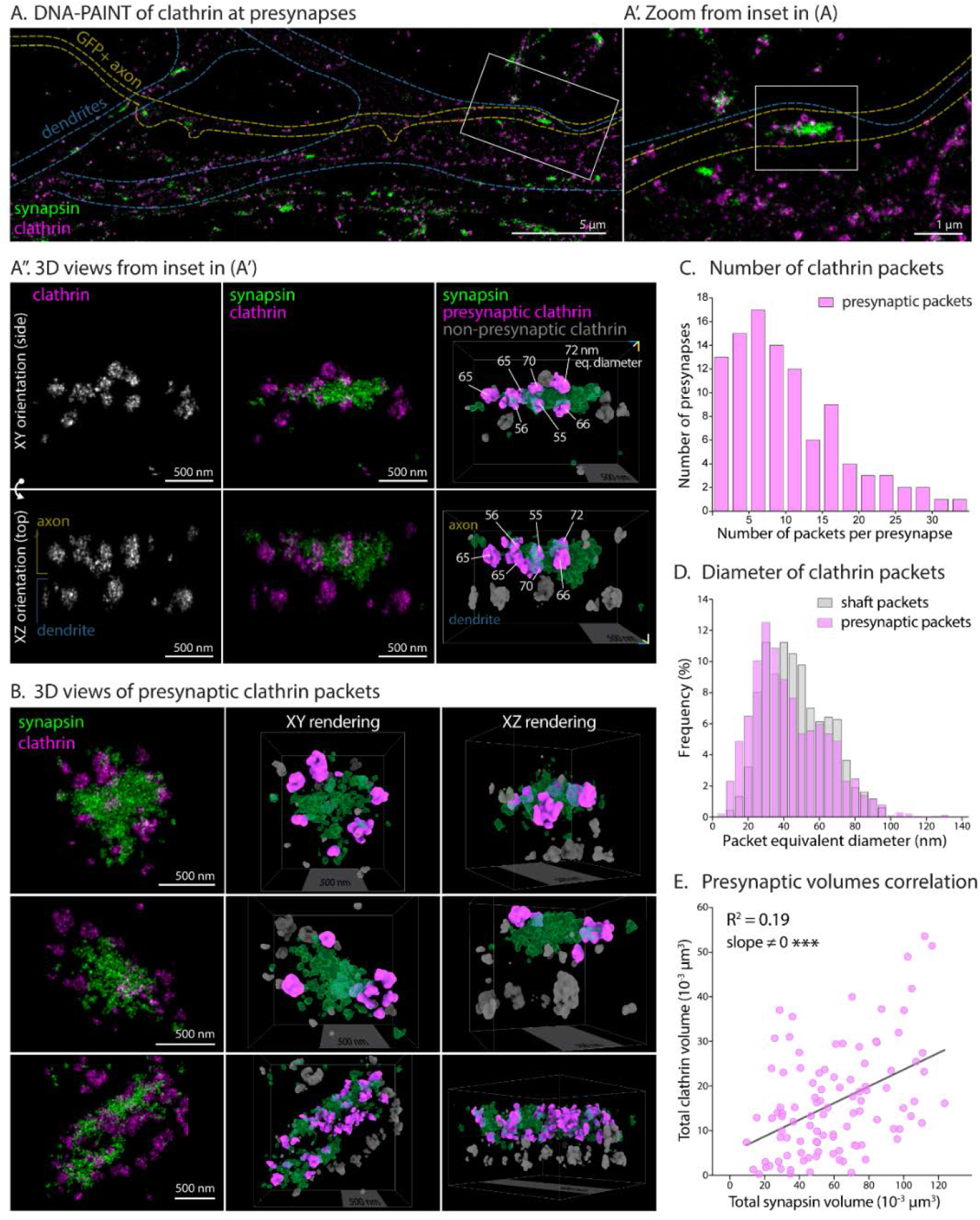
The nanostructure of clathrin at the presynapses. (A) DNA-PAINT of endogenous clathrin (magenta) and synapsin (pseudocolored green, synaptic-vesicle marker). Outlines of dendrites (blue dashed line) and an actin-GFP transfected axon (yellow) are shown; note synapses between GFP+ axon and a large dendrite. (A’) Zoomed view of the white rectangular area in A, centering on a presynaptic bouton and underlying dendrite (note that both axonal and dendritic clathrin particles are seen in this XY view). (A’’) 3-D views of the rectangular inset in (A’), with XY and XZ (rotated) views; along with 3-D rendering and segmentation (right panels). The rendered view shows presynaptic clathrin puncta in magenta, non-presynaptic clathrin in grey, and synapsin in semi-transparent green. Note clathrin particles abutting the presynaptic vesicle cluster. Numbers (white labels) show equivalent diameters calculated from the 3-D renderings. (B) 3D views of additional presynapses showing the XY projection of the DNA-PAINT image (left column, synapsin in green, clathrin in magenta), as well as the XY and XZ view of the segmented rendering (presynaptic clathrin packets in magenta, non-presynaptic clathrin in grey and synapsin in semi-transparent green). (C) Histogram of segmented presynaptic clathrin particles per presynapse (1261 clathrin particles from 102 synapses). (D) Overlaid histograms showing frequency distribution of presynaptic clathrin particle-diameters (n=1261 particles from 3 independent experiments), compared to clathrin particle-diameters in axon shaft (n=685 from 3 independent experiments). (E) The total volume of clathrin (cumulative of all clathrin particles) at a given bouton correlated with the total volume of synapsin.

## DISCUSSION

Using live-imaging, MUDPIT-MS, Apex-labeling with EM-tomography, and 3-D super-resolution/DNA-PAINT, our experiments uncover new structural and molecular aspects of clathrin assemblies in axons and synapses – revealing mechanistic determinants of its slow axonal transport and presynaptic targeting. Axonal clathrin assemblies (transport-packets) move infrequently and intermittently with an anterograde bias, generating an overall biased flow of clathrin in axons. Although clathrin is well-studied in the context of endocytosis, the behavior of axonal clathrin transport-packets seems unrelated to endocytosis, and they appear to be specialized organelles dedicated for transport and targeting. Discrete units of motile clathrin particles resembling transport-packets are seen at presynapses and flanking axons, suggesting targeting and trapping of transport-packets at synapses. Finally, super-resolution-imaging reveals a surprising organization of clathrin at synapses, with multiple clathrin-packets abutting the synaptic-vesicle cluster.

### Clathrin Transport-Packets – a novel Carrier for Slow Axonal Transport

Mechanisms of slow axonal transport – cargoes moving for days to months along axons – are intriguing. A better understanding of cargoes moving in slow transport might bring clarity, just like visualization of vesicle-transport in the 1980’s triggered insights into vesicle transport – including the discovery of kinesins (Grafstein and Forman, 1980; Maday et al., 2014). Though pulse-chase radiolabeling studies have established that clathrin is conveyed in the slow component, the methods used could not visualize the movement, and our original motivation was to determine the cargo-structure moving clathrin in slow axonal transport. Given the known transient nature of clathrin in non-neuronal cells and dendrites, we were surprised to see long-lasting and motile axonal clathrin particles that are unrelated to endocytosis. Though previous studies in non-neuronal cells have described motile clathrin-coated intermediates in the cytoplasm (Keyel et al., 2004; Puertollano et al., 2003; Rappoport and Simon, 2003; Rappoport et al., 2003a; Rappoport et al., 2003b), these seem different from the axonal transport-packets. First, the clathrin intermediates in non-neuronal cells are large, pleomorphic tubulovesicular aggregates (Polishchuk et al., 2006; Puertollano et al., 2003); different from the uniform-sized structures we see in the axon (see ultrastructure in **Fig. 4** and **Supp. Movies 3 and 4**). Though clathrin intermediates in non-neuronal cells can move, the clathrin is reported to “cycle on and off” – eventually fusing with early or late endosomes (Puertollano et al., 2003) – which is also different from what we see in axons. Thus, we posit that neurons may have evolved specialized mechanisms for biased slow transport of clathrin.

The motility of the clathrin transport-packets in axons is microtubule-dependent (**Fig. 3D-E**). Previous studies have shown associations between tubulin and clathrin in non-neuronal cells (Kelly et al., 1983; Pfeffer et al., 1983). Since the instantaneous velocities of the packets resemble motor-driven cargoes, motor proteins are likely involved (dynein heavy chain and myosin-10 were seen in our MS data, besides myosin-6; see **Supp. Table 1**). Whether the motor proteins directly bind to clathrin, or some other component of the transport-packet remains to be determined, but its intriguing to note the close apposition of the clathrin “spokes” with microtubules in our EM-tomograms (see **Supp. Movie 4)**. Our proteomics data also showed an association of clathrin with myosin-6; and further experiments suggested that the latter plays a role in anchoring clathrin particles in axons (**Figs. 5–6**). The close apposition of clathrin transport-packets and the sub-membranous actin-spectrin cytoskeleton in axons (see **Fig. 4C**) also points to a link between transport-packets and actin. Although disruption of actin/myosin-6 did not eliminate the stationary clathrin particles in axons and it is likely that there are other players involved in anchoring axonal clathrin; to our knowledge this is the first insight into mechanisms that might act as physiologic tethers in any form of slow transport.

### A diversity of mechanisms in Slow Axonal Transport

Cytosolic proteins moving in slow axonal transport are surprisingly diverse – proteins involved in metabolism, synaptic homeostasis, cytoskeletal proteins, endocytosis-related proteins, etc. Based on the apparently coherent-migration of diverse cytosolic proteins in radiolabeling experiments, a prevailing notion in the field is that there is a unified mechanism moving all cytosolic proteins in slow axonal transport – sometimes called the “structural hypothesis” [(Garner and Lasek, 1982; Lasek et al., 1984; Tytell et al., 1981) – reviewed in (Black, 2016; Roy, 2014)]. However, our studies in cultured neurons, visualizing and manipulating cytosolic cargoes at high resolution, suggest that mechanistic perturbations of one cargo has no effect on the dynamics of another. For instance, though inhibition of myosin-6 accelerates the dynamics of clathrin transport-packets, it has little effect on the anterogradely-biased transit of the slow-component protein synapsin (**Supp. Fig. 5C-D**). Analogously, Hsc70-inhibition – that blocks the anterogradely-biased mobility of synapsin in axons, as shown in our prior study (Ganguly et al., 2017) – has no effect on axonal clathrin dynamics (**Supp. Fig. 5E**). Moreover, the overall morphology and transport-behavior of these two slow-component cargoes is very different. While synapsin and dynein are homogenously distributed in axons, and the anterograde transport is likely generated by a “biased flow” (Scott et al., 2011; Tang et al., 2013; Twelvetrees et al., 2016); the axonal distribution of clathrin is clearly particulate, and the anterograde bias is generated by the motility of discrete particles (see **Fig. 2**). There are also significant differences in our proteomics interactome-data from synapsin- and clathrin-associated proteins; though the starting-fractions were not the same in the two studies [(Ganguly et al., 2017) and see **Fig. 5**]. The slow transport of cytoskeletal cargoes seems to employ yet different mechanisms (Chakrabarty et al., 2019; Roy et al., 2000; Wang et al., 2000); and its highly unlikely that the structural hypothesis, as originally stated, is a valid model of slow transport. Nevertheless, an emerging theme is that cohorts of functionally related cytosolic proteins – for instance “synaptic” or “endocytic” groups – might be co-transported as multiprotein complexes, so they can function together; a topic for future studies.

### Dynamics and Organization of Clathrin at presynapses

The distribution of clathrin at synapses was not homogenous but appeared as multiple motile puncta, moving within individual boutons and flanking axons (**Fig. 7**). Clathrin puncta frequently moved between multiple adjacent boutons (**Fig. 7**), reminiscent of mobile synaptic-like vesicles that also do the same (Darcy et al., 2006). Synapses contained a discrete number of clathrin-assemblies resembling transport-packets, suggesting that the latter were trapped and retained at boutons. Our DNA-PAINT data – staining endogenous synaptic proteins – reveal a striking pattern with multiple clathrin puncta around the synaptic-vesicle cluster, with clathrin particles extending into flanking axons (**Fig. 8**). Though previous EM studies of synapses have seen intact clathrin-coated vesicles (Tao-Cheng, 2020), the abundance and organization of clathrin-assemblies seen in our experiments is surprising. Presumably the 3-D views, with high-resolution and signal-to-noise ratio of DNA-PAINT allowed us to pinpoint the clathrin-packets, which may have been hard to discern in conventional 2D EM from thin-sections. Finally, it seems likely that the radial organization of clathrin around synaptic-vesicle clusters has functional implications. Indeed in addition to the classic model of clathrin-mediated endocytic retrieval of synaptic vesicles (Heuser and Reese, 1973), clathrin-independent modes are being increasingly recognized as important players (Chanaday and Kavalali, 2018; Farsi et al., 2018; Soykan et al., 2017; Watanabe et al., 2013); and perhaps a better understanding of the spatial organization of endocytic proteins will bring some clarity.

## Acknowledgements

This work was supported by NIH grants to SR (R01NS075233, R01AG048218); a CNRS ATIP grant to CL (ATIP2016); and NIH grants to DB (R01GM086197) and MHE (P41 GM103412), for support of the National Center for Microscopy and Imaging Research. The authors thank Leonardo Parra for helpful comments on the manuscript, and various investigators for generously sharing constructs (listed in the methods section).

## METHODS

### DNA constructs, pharmacologic agents, neuronal cultures, and transfections

The GFP:CLC, pUC57-APEX and Dendra2 constructs were obtained from Addgene (from the laboratories of Drs. Peter McPherson [McGill University, Canada], Alice Ting [Stanford University, CA] and Michael Davidson [Florida State University, Tallahassee, FL], respectively). The PAGFP:CLC construct was subcloned from the PAGFP backbone, a gift from Dr. Jennifer Lippincott-Schwartz (Janelia Farms Research Campus, Ashburn VA) by standard cloning. CHC-T7-Hub and T7-empty backbone was a gift from Dr. Frances Brodsky (UCL, London, UK). The GFP:Utr-CH, NPYss:mCherry, and Synaptophysin:DsRed constructs were gifts from Drs. William Bement (University of Wisconsin, Madison, WI), Gary Banker (Oregon Health & Science University, Portland, OR) and Leon Lagnado (University of Sussex, Sussex, England, UK) respectively. The MyoVI-DN and the MyoVI full length constructs were a gift from Dr. Don Arnold (University of Southern California, Los Angeles CA). Dynasore and Nocodazole were dissolved in DMSO and used at a final concentration of 80 μM and 10 μg/ml. Swinholide A were dissolved in ethanol and used at a final concentration of 100 nM respectively. All drugs were purchased from Millipore-Sigma and added to Hibernate E media (see below) prior to imaging.

All hippocampal cultures were obtained from brains of postnatal (P0–P1) CD-1 mice and plated on MatTek glassbottom dishes as described previously in detailed published protocols (Ganguly and Roy, 2014; Roy et al., 2012), in accordance with University of California guidelines. In brief, MatTek dishes were coated with 100 μl of 1 mg/ml poly-d-lysine for 2 h at RT, washed thrice with ddH2O, and air dried before plating. Hippocampi from P0–P1pups were dissected in ice-cold dissection buffer (HBSS, 4.44 mM d-glucose, and 6.98 mM HEPES) and incubated in 0.25% Trypsin–EDTA at 37°C for 15 min. Following this, neurons were dissociated in plating media (10% FBS and 90% Neurobasal/B27; Life Technologies) by trituration. Neurons were plated at a density of 50,000 cells/100 μl (for clathrin imaging at en passant boutons) and at 25,000 cells/100 μl (for all other experiments) of plating media. Neurons were maintained in Neurobasal/B27 media (supplemented with 2% B27 and 1% GlutaMAX) in an incubator at 37°C and 5% CO_2_ for 7–9 d before transfection. Neurons were transfected with the indicated fluorescent constructs (0.8 μg DNA for clathrin constructs, 1.2 μg of all others) at 7–9 DIV with Lipofectamine 2000 (Life Technologies), and all live imaging was performed between 12–16 h after transfection. Neurons were co-transfected with soluble GFP or mCherry to identify axons and dendrites. GFP:CLC transfected neurons selected for imaging had punctate clathrin appearance in dendrites, and axon regions imaged were more than 150 μm from the neuronal cell body.

### Live imaging of axons and synapses

Most single-color live imaging experiments were performed on an Olympus IX81 inverted motorized epifluorescence microscope equipped with a CoolSNAP *HQ2* camera (Photometrics). Details of the photoactivation and photoconversion setup have been described previously (Roy et al., 2012). Briefly, a dual-source light illuminator (IX2-RFAW, Olympus) was used to simultaneously visualize and photoactivate/photoconvert a discrete ROI within the axon. Before live imaging, neurons were transferred to Hibernate media (Brainbits), supplemented with 2% B27, 2mM Glutamax, 0.4% D-glucose, 37.5 mM NaCl (Ganguly et al., 2017; Roy et al., 2012; Scott et al., 2011), and maintained at 37°C (Precision Control Weatherstation) for the duration of the experiments. Axons were identified based on morphology, and only neurons with unambiguous morphology were selected for imaging (Roy et al., 2012; Scott et al., 2011). For GFP:CLC, neurons were imaged every 0.7 s for several minutes. For photoactivation experiments, PAGFP:CLC was photoactivated with 405 nm light for 1 s, and imaged every 1.2 s. Dendra2:CLC was photoconverted by 405 nm light for 3 s, and imaged every 1.2 s. Synaptophysin:GFP and NPYss:mCherry was imaged using the stream acquisition function (MetaMorph) at 5 frames/s (with no time interval between images). For imaging *en passant* boutons, 9–12 DIV neurons were co-transfected with synaptophysin:dsRed and GFP:CLC, and boutons were identified based on size of the synaptic vesicle cluster and morphology (Wang et al., 2014). Boutons along the primary axon, >150 μm from the soma were selected for imaging. Additionally, axons selected for imaging had at least two or more en passant boutons in proximity and imaged at 1 s interval. Dual-color live imaging of mCherry:CLC and GFP:Utr-CH was done on an inverted epifluorescence microscope (Eclipse Ti-E; Nikon) equipped with CFI Plan Apochromat VC 100× oil (NA 1.40; Nikon) objective, and an electron-multiplying charge-coupled device camera (QuantEM:512SC; Photometrics). Exciting red/green LED lights were rapidly switched (within microseconds) using the SPECTRA X LED illuminator, achieving near-simultaneous two-color imaging. A multi– band-pass filter (Chroma Technology Corp.) was inserted into the emission light path, and GFP/RFP images were obtained with precise subpixel registration. All movies were analyzed using kymographs generated with a built-in function in MetaMorph (Molecular Devices, LLC). Photoactivation and photoconversion data was analyzed using the intensity-center/bin-center shift method described previously in several publications from our group (Chakrabarty et al., 2019; Ganguly et al., 2017; Roy et al., 2012; Scott et al., 2011; Tang et al., 2013). All statistical analysis was performed using Graph Pad Prism (Graph Pad Software, San Diego, CA).

### Biochemical fractionation, western blotting and immuno-precipitation from mouse sciatic nerves

In vivo biochemical assays were adapted from protocols described earlier by (Das et al., 2013; Ganguly et al., 2017; Scott et al., 2011; Tang et al., 2012). Briefly, sciatic nerves were dissected from 6-8-wk-old mouse CD1 (WT) mouse. 64 sciatic nerves were pooled for each round of immune-precipitation (IP) and crushed in liquid nitrogen using a motor pestle and then homogenized in nondenaturing buffer (1X IP buffer, Invitrogen Dynabeads™ Co-Immunoprecipitation Kit [ThermoFisher Scientific, USA]) using 18 G and 23 G needles in the presence of protease inhibitor cocktail (Sigma-Aldrich). The resulting homogenate was centrifuged at 1,000 g for 20 min at 4°C to obtain a nuclear pellet (P1) and a post-nuclear supernatant (S1). The S1 supernatant was then centrifuged at 10,200 g for 20 min at 4°C to obtain a crude synaptosomal fraction (P2) and synaptosome-depleted fraction (S2). IP was performed using the DynaBeads Co-Immunoprecipitation kit (14321D; Thermo Fisher Scientific). After centrifugation, the S2 fraction was divided equally and incubated with 10.5 mg anti–Clathrin-Heavy-Chain antibody (ab21679; Abcam, Cambridge MA) and anti-Mouse IgG (ab190475; Abcam, Cambridge MA) for overnight coupling with magnetic beads at 37°C. All following washes were performed as per the manufacturer’s protocol. The S2 fraction was incubated with the antibody-coupled beads for 35 minutes at 4°C on a rotor. After the final wash, 3 mg beads from each fraction were subjected to 2D gel electrophoresis on a 4-12% gradient SDS-page gel and then probed with the anti–Clathrin-Heavy-Chain antibody to determine the efficacy of immunoprecipitation. The remaining 7.5 mg of S2 fraction lysate beads were then used for MudPIT-MS analysis. Two independent repeats were performed with sciatic nerve axons S2 lysate.

### Protein identification through MudPIT-MS analysis and in silico data analysis

MudPIT-MS protocols were identical to our previous studies (Ganguly et al., 2017). Briefly, beads coated with the clathrin heavy chain antibody (or mouse IgG) were dissolved in 100 μl of 8 M urea in 100 mM Tris, pH 8.5, followed by reduction and alkylation in 10 mM Tris (2-carboxyethyl) phosphine hydrochloride (Roche) and 55 mM iodo-acetamide (Sigma-Aldrich), respectively. This was followed by digestion in trypsin (Promega; incubated at 37°C overnight in the dark) and magnetic bead removal by a magnetic separator. The resulting protein digest was acidified in formic acid followed by centrifugation at 14,000 rpm for 10 min. Thereafter, the supernatant was pressure loaded onto a 250 μm inner diameter– fused silica capillary column (Polymicro Technologies). This column was fitted with a Kasil frit packed with 2.5 cm of 5 μm Partisphere strong cation exchange resin (Whatman) and 2.5 cm of 5 μm C18 resin (Phenomenex). After desalting, this biphasic column was connected to a 100 μm inner diameter-fused silica capillary (Polymicro Technologies) analytical column with a 3 μm pulled tip packed with 10 cm of 3 μm C18 resin (Phenomenex). The entire three-phase column was then laced in line with a 1,200 quaternary HPLC pump (Agilent Technologies) and analyzed using a modified 12-step separation described previously (Washburn et al., 2001). As peptides were eluted from the microcapillary column, they were electrosprayed directly into a hybrid LTQ Orbitrap Velos mass spectrometer (Thermo Fisher Scientific). A cycle consisted of one full-scan mass spectrum (300–1,600 m/z) followed by 20 data-dependent collision-induced dissociation tandem MS spectra. The application of mass spectrometer scan functions and HPLC solvent gradients was controlled by the Xcalibur data system (Thermo Fisher Scientific). Tandem MS spectra were extracted using RawXtract (version 1.9.9; (McDonald et al., 2004)) and searched with the ProLuCID algorithm (Xu et al., 2015) against a mouse UniProt database concatenated to a decoy database in which the sequence for each entry in the original database was reversed (Peng et al., 2004). The ProLuCID search was performed using semienzyme specificity and static modification of cysteine because of carboxyam-idomethylation (57.02146). ProLuCID search results were assembled and filtered using the DTA Select algorithm (version 2.0; (Tabb et al., 2002)). The protein identification false positive rate was kept below 1%, and all peptide– spectra matches had <10 ppm mass error.

For selection of peptides from the raw MudPIT data, peptides were subjected to the following selection criteria. First, the same peptide fragment had to appear in both rounds of the co-immunoprecipitation, followed by the criteria that the total raw spectrum count from both rounds for the peptide should be >30. Finally, only peptides with combined spectrum counts twenty-fold or higher than the combined spectrum counts from IgG coupled beads were selected. Only peptides that meet all three criteria were included in the final list (**Supp. Table 1**). The identification of peptides and gene names was performed using the Uniprot database (https://www.uniprot.org/). For generation of the clathrin protein interaction map from the S2 fraction all known interactions from experimental data were determined from the String protein interaction database (http://www.string-db.org).

### Sample preparation, data acquisition and data processing for electron tomography

For EM, mouse hippocampal neurons were plated at a density of 25,000 cells/100 μl in wells of gridded MatTek dishes coated with poly-D-lysine. DIV 9-10 neurons were transfected with Apex:GFP:CLC using lipofectamine 2000 as described previously (Ganguly et al., 2017). 6-8 hours before transfection, 7 μM Hemin chloride (Sigma-Aldrich) was added to the culture medium and kept in the medium until the fixation step (below). 12-16 hours after transfection, transfected neurons were identified based on the GFP signal, and corresponding grid numbers were noted. Following this, neurons were fixed for 5 mins at room temperature, and then 60 minutes on ice in 2.5 % glutaraldehyde in 0.1M sodium cacodylate buffer (pH 7.4). After washes in cacodylate buffer and a quenching step in 20mM glycine, cells were incubated with freshly prepared DAB (25.24 mM) and 0.03% H_2_O_2_ for 25-30 minutes. Neurons were then washed in 0.1M cacodylate buffer and post-fixed with 1% osmium tetroxide for 30 minutes on ice. The neurons were washed with ddH2O and then dehydrated and embedded in Durcupan epoxy resin. After curing, the coverslip from the bottom of the MatTek dish was removed gently, and grid numbers with transfected neurons were sawed out and mounted on a dummy acrylic block for sectioning. Ultra-thin (70 nm) and semi-thin sections (200 nm) were cut using a diamond knife (Diatome). To improve stability of specimens under the beam of the EM for tomography, these sections were coated with carbon on both sides. Colloidal gold particles (5 nm and 10 nm diameter) were deposited on each side of the sections to serve as fiducial markers.

Ultra-thin sections were imaged on the JOEL 1200V while the EM tomogram data were obtained from distal axonal regions (>150 μm from the cell body) of transfected neurons using a FEI Titan high base microscope operated at 300 kV and micrographs were produced using a 4K × 4K Gatan CCD camera (US4000). Both microscope and detector were controlled by the Serial EM software package (Mastronarde, 2005) which managed the automated tilt series acquisition. Technical details of how the sections of the tomogram were acquired, image series were aligned properly, and its 3D representations were created have been described by (Phan et al., 2017). Briefly, sections were tilted every 1° from −60° to +60°, aligned properly using colloidal gold particles as fiducial markers and final high-quality 3D representations were built from the projection sets using a custom written non-linear bundle adjustment scheme in TxBR (Lawrence et al., 2006). 3D tomograms were visualized in IMOD (Kremer et al., 1996) and only axon regions with clathrin organelles which do not open to the surface were selected for representative images.

### DNA-PAINT imaging and image analysis

Rat hippocampal neurons were cultured on 18 mm coverslips at a density of 6,000/cm from embryonic day 18 pups following established guidelines of the French Animal Care and Use Committee (French Law 2013-118 of 1st February 2013) and approval of the local ethics committee (agreement 2019041114431531-V2 #20242). To selectively label identified axons, neurons in some experiments were transfected the day before fixation with actin-GFP using Lipofectamine 2000 according to the manufacturer’s instructions. After 9 to 13 DIV, neurons were fixed at RT in phosphate buffer (PB) 0.1M pH 7.3 containing 4% paraformaldehyde and 4% sucrose for 10 minutes, followed by permeabilization and blocking in incubation buffer (IB): 0.22% gelatin, 0.1% Triton in PB 0.1M pH 7.3 at RT for 2 hours, and incubated overnight at 4°C with primary antibody in IB: polyclonal chicken anti-Map2 (Abcam #53392, 1:1000), polyclonal rabbit anti-clathrin heavy chain (Abcam #21679, 1:150) and monoclonal mouse anti β2-spectrin (for axon shaft images, BD Bioscience 612563, 1:150) or monoclonal mouse anti-synapsin (Synaptic Systems #106-001, 1:500. After rinses, neurons were incubated with secondary antibodies, including anti-rabbit D2 and anti-mouse D2 DNA-coupled secondary antibodies for DNA-PAINT (Ultivue) at RT for 1 hour.

DNA-PAINT imaging (Jungmann et al., 2014) was performed on an N-STORM microscope (Nikon Instruments) in imaging buffer with 0.25-1 nM Imager-650 and nM Imager-560 (Ultivue). The sample was alternatively illuminated at 647 nm and 561 nm (full laser power) and 20,000-30,000 images of each channel were acquired at 25-33 Hz. DNA-PAINT acquisitions were processed using the N-STORM software followed by post-processing; filtering and image reconstruction at 4 or 12 nm/pixel with the ThunderSTORM Fiji plugin (Jimenez et al., 2019). For 3D rendering, localizations were density-filtered (50 nm 3D radius) before reconstruction of individual presynapses as of 4-nm voxel stacks. Stacks where then 3D gaussian-smoothened and imported into ChimeraX software (Goddard et al., 2018). After surface thresholding and smoothing (default values), 3D rendering images and movies were exported as images and movies. Clathrin particles were segmented using the Segger plugin and their volume measured. Volume V in μm3 was converted in equivalent sphere diameter D in nm using [D=1000*2*V(^1/3)*3/(4*π)] for graphing. For presynaptic clathrin particles, only particles contacting or closely apposed to the synapsin cluster were counted and measured.

## SUPPLEMENTARY FIGURES

**Supplementary Figure 1:**
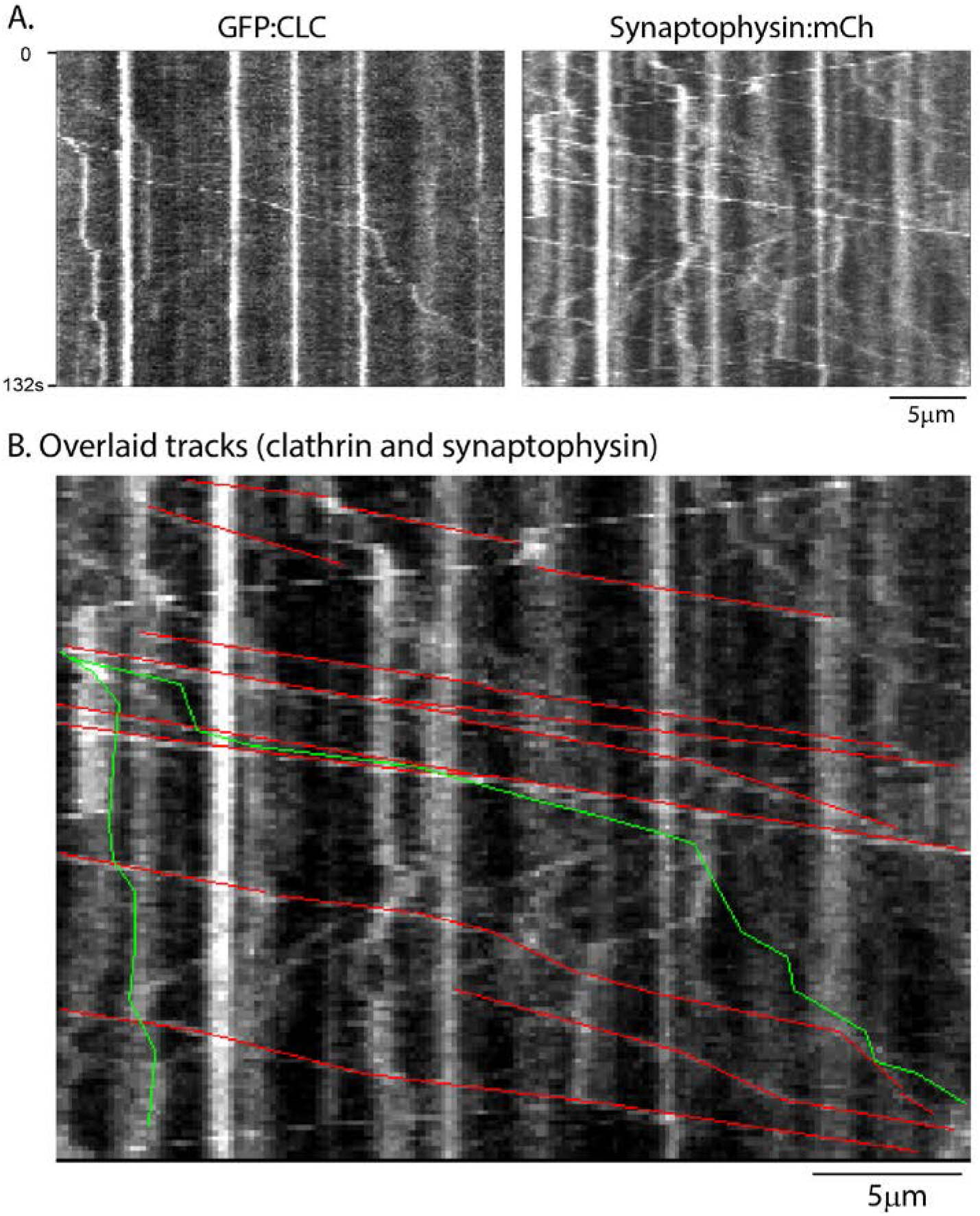
Near-simultaneous imaging of clathrin transport-packets and vesicles. Kymographs from dual-imaging of GFP:CLC and synaptophysin:mCherry (top), with overlaid tracks (bottom). The movement of clathrin transport-packets was clearly much less frequent than the fast transport of synaptophysin-vesicles. Though the extensive movement of synaptophysin made definitive conclusions difficult, we could see clear instances where clathrin-packets were transported independently of synaptophysin – see overlaid clathrin (green) and synaptophysin (red) tracks in kymograph below.

**Supplementary Figure 2:**
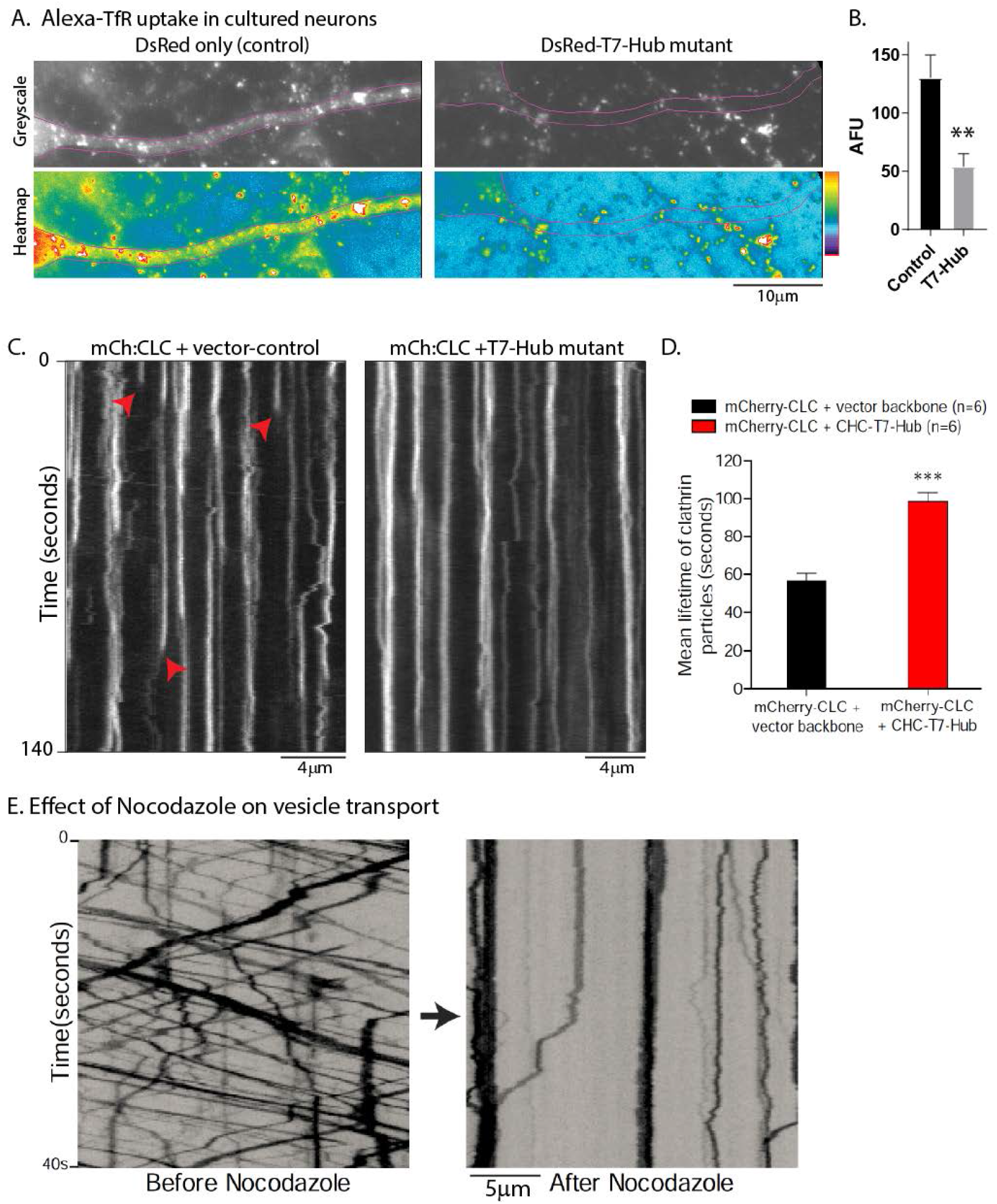
The clathrin T7-Hub mutant blocks receptor mediated endocytosis. (A) Neurons were transfected with T7-Hub mutant (DsRed-tagged) or vector-control (DsRed-only), and Alexa-488-transferrin uptake into the transfected neurons was analyzed. As evident in the representative greyscale images (above) and pseudocolor heatmaps (below), the T7-Hub mutant attenuates transferrin-uptake. Data quantified in (B), 5-6 neurons per condition, from one culture; **p<0.001. (C) Kymographs showing dynamics of mCh:CLC in dendrites co-transfected with the T7-Hub mutant (or vector control). Note that the blinking of clathrin fluorescence – due to receptor mediated endocytosis, some marked by arrowheads – is essentially abolished in the T7-Hub transfected neurons. (D) Quantification of mCh:CLC lifetime in dendrites (100-150 particles from 6 dendrites were analyzed; ***p<0.0001). (E) Kymographs showing movement of the pan-vesicle marker NPYss:mCherry (see Ganguly et al., 2015) in an axon before and after 10μg/ml Nocodazole for 30 min. Note complete stoppage of vesicle transport after adding the microtubule depolymerizing agent.

**Supplementary Figure 3:**
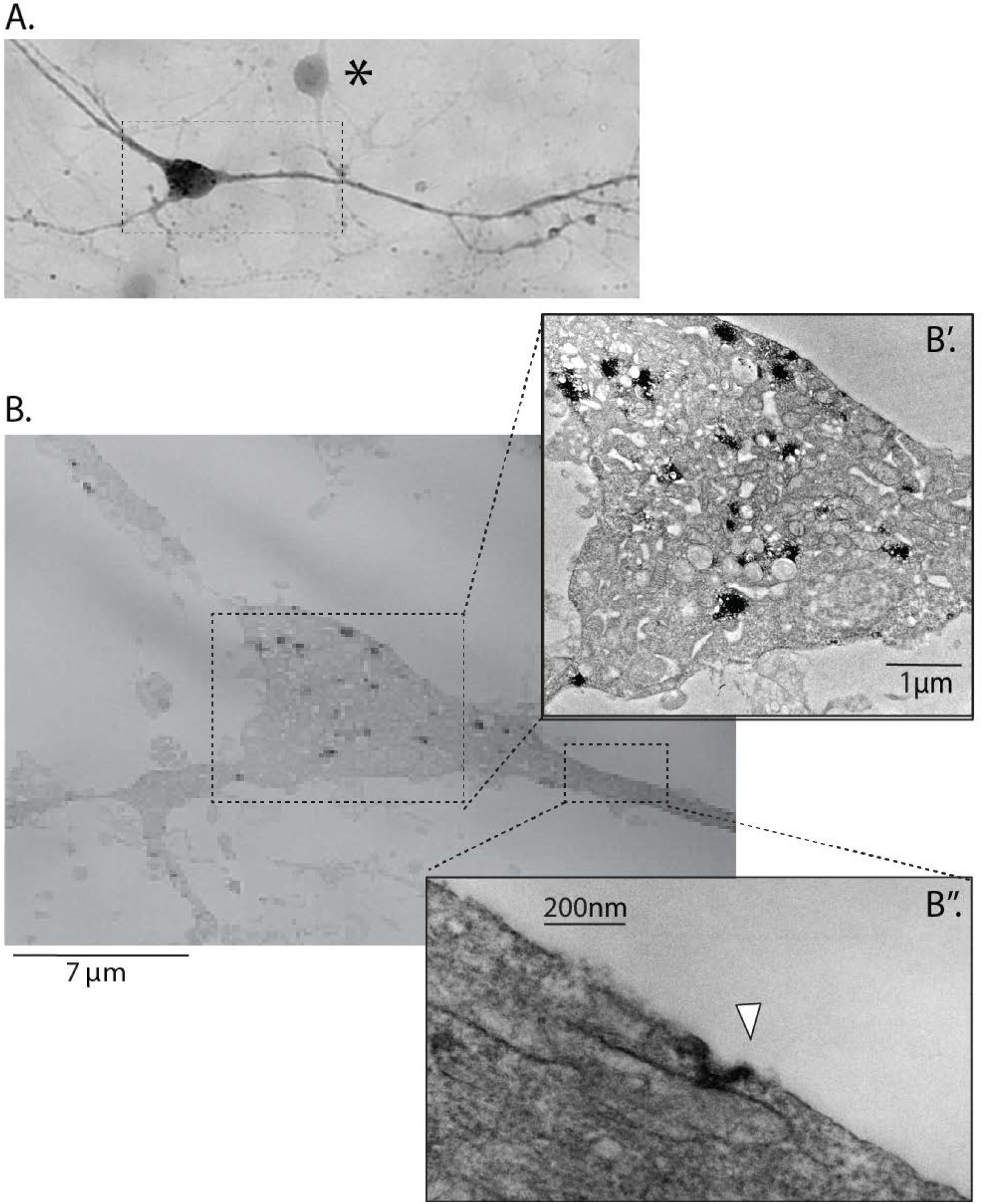
Apex-clathrin labeling and EM. (A) Neurons were transfected with Apex:GFP:CLC and processed for EM. Gridded coverslips were used to identify transfected neurons for EM processing, as described in methods. The DAB signal from an Apex:GFP:CLC transfected neuron is shown, along with an adjacent un-transfected neuron (marked with an asterisk). (B) EM of rectangular region from (A), dark signals represent clathrin. Note that many clathrin structures are seen in the somatodendritic region (B’), some clearly representing coated clathrin pits (B’’).

**Supplementary Figure 4:**
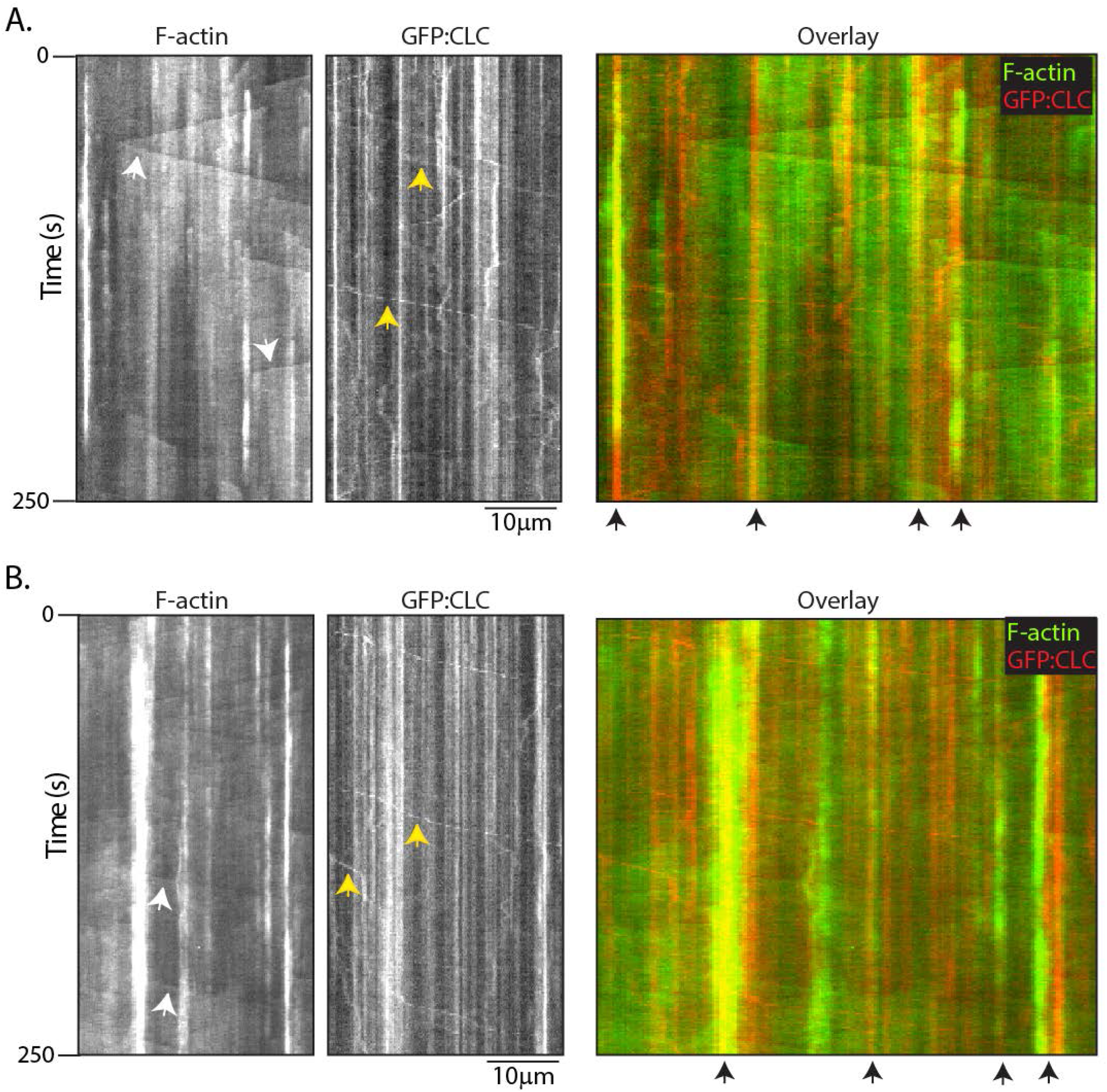
Dual-color live imaging of F-actin and clathrin. Kymographs from two-color imaging of GFP:Utr-CH (F-actin) and mCh:CLC – some ‘actin trails’ and clathrin transport-packets are marked by white and yellow arrowheads respectively. On the overlaid red/green kymographs on right, note that many stationary clathrin particles are colocalized with F-actin (black arrowheads, bottom).

**Supplementary Figure 5:**
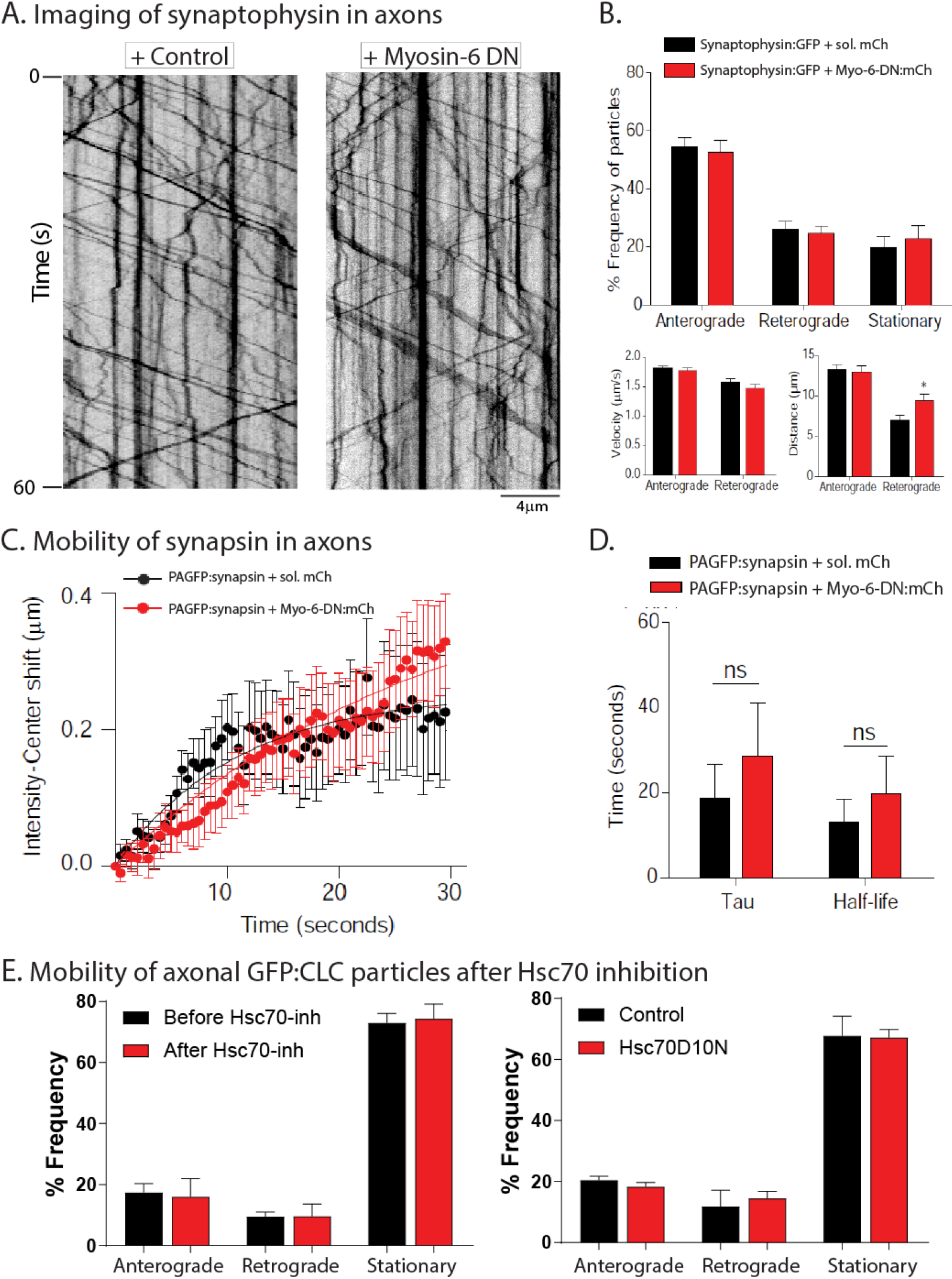
Disruption of Myosin-6 has no effect on fast synaptophysin or slow synapsin transport. (A) Kymographs of synaptophysin:GFP transport in neurons transfected with dominant-negative Myosin-6 (or control vector). Note that disruption of Myosin-6 has no effect on the fast transport of synaptophysin-labeled vesicles. Data quantified in (B). (~ 350-450 mobile particles from ~ 10 axons were analyzed; data from three separate cultures). (C) Ensemble data of axonal intensity-center shifts from neurons transfected with PAGFP:synapsin (slow transport assay). Neurons were also co-transfected with dominant-negative Myosin-6 (or control vector). Note that disruption of Myosin-6 has no significant effect on the anterograde bias of synapsin in axons; quantified in (D). (15 neurons from three separate cultures were analyzed for each group). (E) Quantification of GFP:CLC particle dynamics in axons before and after pharmacologic inhibition of Hsc70 with a small-molecule (VER155008); or genetic interference of Hsc70 function using a dominant-negative construct (Hsc70D10N). Note that motility of clathrin is unchanged after Hsc70 inhibition.

## SUPPLEMENTARY MOVIE LEGENDS

**Supplementary Movie 1:** GFP:CLC dynamics in dendrites, movie corresponds to kymograph shown in **Figure 1B**. Short vertical lines mark regions where clathrin puncta appear and disappear, indicating clathrin mediated endocytosis. Time in seconds is on upper left and scale bar is 5 μm.

**Supplementary Movie 2:** GFP:CLC dynamics in axons, movie corresponds to kymograph shown in **Figure 1C**. Note rapid vectorial movement of clathrin particles. Time in seconds is on lower left and scale bar is 5 μm.

**Supplementary Movie 3:** EM tomogram of Apex2-tagged clathrin in an axon-shaft (contrast-enhanced to show details; see methods for correlative light/EM); **Figure 4B”** is a still frame from this movie. Red arrows point to three axonal clathrin coated structures, presumably representing transport-packets. In the 3-D views, note proximity of the clathrin structures to microtubules running along the longitudinal axis (also see **Supp. Movie 4**). Also note that axons contain many other non-clathrin coated vesicles including endolysosomal organelles.

**Supplementary Movie 4:** EM tomogram of Apex2-tagged clathrin in an axon-shaft (contrast-enhanced to show details; see methods for correlative light/EM). Red arrow points to a clathrin coated structure. This 3-D view shows a clear apposition of clathrin with a microtubule, with some of the “spokes” appearing to touch the microtubules.

**Supplementary Movie 5:** 3-D rendering and segmentation from DNA-PAINT super-resolution images of an axon stained with β2-spectrin and clathrin heavy-chain antibodies (corresponds to **Figure 4C**). β2-spectrin – used here as a marker of axonal boundary – is in semi-transparent green, and magenta marks axonal clathrin. Some very small particles are rendered grey. Note many clathrin structures resembling transport-packets, often seen in close apposition to the sub-plasmalemmal β2-spectrin lattice (see results).

**Supplementary Movie 6:** GFP:GFP:CLC dynamics in *en-passant* presynaptic boutons (identified by synaptophysin:mCherry, see results). Movie corresponds to still-frames shown in **Fig. 7A**. Note multiple motile puncta within boutons – resembling transport packets seen in the axon-shafts – and intra-bouton exchange of clathrin particles. Time in seconds is on lower left and scale bar is 5 μm.

**Supplementary Movie 7:** 3-D rendering and segmentation from DNA-PAINT super-resolution images of four presynaptic boutons stained with clathrin heavy-chain and synapsin antibodies – the latter to mark the synaptic-vesicle cluster (movie corresponds to **Figure 8**). Clathrin is in magenta, and synapsin is in semi-transparent green. Non-synaptic clathrin particles are shown in grey. Note multiple clathrin particles abutting the synaptic-vesicle cluster.

