## Supplementary material for "Mechanistic Determinants of Slow Axonal Transport and Presynaptic Targeting of Clathrin Packets": Supp. Table 1

| Protein Families | UniProtKB ID (from MudPIT) | Gene IDs | Organism |
|---|---|---|---|
| ATP citrate synthase | Q3V117 | Acly | Mus musculus (Mouse) |
| Actin | P68134, P60710, Q8BFZ3, P68033 | Acta1, Actb, Actbl2, Actc1 | Mus musculus (Mouse) |
| Argonaute | Q8CJG0 | Ago2 | Mus musculus (Mouse) |
| Ahnak2 | E9PYB0 | Ahnak2 | Mus musculus (Mouse) |
| Adaptor Related Protein Complex | Q8CC13, Q8CBB7, Q3UHJ0, P17426, P17427, Q9DBG3, P84091, P62743, Q9Z1T1 | Ap1b1, Ap1g1, Aak1, Ap2a1, Ap2a2, Ap2b1, Ap2m1, Ap2s1 , Ap3b1 | Mus musculus (Mouse) |
| V-ATPase | Q9Z1G4 | Atp6v0a1 | Mus musculus (Mouse) |
| Bag3 | Q9JLV1 | Bag3 | Mus musculus (Mouse) |
| Bcr | Q6PAJ1 | Bcr | Mus musculus (Mouse) |
| Bmp | Q91Z96 | Bmp2k | Mus musculus (Mouse) |
| Calcoco | Q8CGU1 | Calcoco1 | Mus musculus (Mouse) |
| Ccdc | D3YZP9 | Ccdc6 | Mus musculus (Mouse) |
| Clint1 | Q5SUH7 | Clint1 | Mus musculus (Mouse) |
| Clathrin | Q6IRU5, Q68FD5 | Cltb, Cltc | Mus musculus (Mouse) |
| Alpha-crystallin | P23927 | Cryab | Mus musculus (Mouse) |
| Casein | P19228, Q02862 | Csn1s1, Csn1s2a | Mus musculus (Mouse) |
| Dab2 | P98078 | Dab2 | Mus musculus (Mouse) |
| Connecdenn | Q8K382 | Dennd1a | Mus musculus (Mouse) |
| Dynamin | P39053, P39054 | Dnm1, Dnm2 | Mus musculus (Mouse) |
| Dynein | Q9JHU4 | Dync1h1 | Mus musculus (Mouse) |
| Edc | Q3UJB9 | Edc4 | Mus musculus (Mouse) |
| Eef | P58252 | Eef2 | Mus musculus (Mouse) |
| Epsin | Q80VP1, Q5NCM5 | Epn1, Epn2 | Mus musculus (Mouse) |
| Eps | P42567, Q60902 | Eps15, Eps15l1 | Mus musculus (Mouse) |
| Fatty acid binding protein | Q05816 | Fabp5 | Mus musculus (Mouse) |
| Fatty Acid Synthase | P19096 | Fasn | Mus musculus (Mouse) |
| Fcho | Q3UQN2 | Fcho2 | Mus musculus (Mouse) |
| Fibrinogen A | E9PV24 | Fga | Mus musculus (Mouse) |
| Filamin A | Q8BTM8 | Flna | Mus musculus (Mouse) |
| Gak | Q99KY4 | Gak | Mus musculus (Mouse) |
| Gulp1 | Q8K2A1 | Gulp1 | Mus musculus (Mouse) |
| H1 | P15864, P43274 | H1-2, H1-4 | Mus musculus (Mouse) |
| Hip1 | Q8VD75, Q6ZQ77 | Hip1, Hip1r | Mus musculus (Mouse) |
| Hnrnp | Q8BG05, Q8VDM6 | Hnrnpa3, Hnrnpul1 | Mus musculus (Mouse) |
| Heat shock proteins | Q80TZ3, Q71LX8, Q8K0U4, Q61696, P16627,P17156, Q3U2G2, P48722, P20029, Q504P4, Q61699 | Dnajc6, Hsp90ab1, Hspa12a, Hspa1a, Hspa1l, Hspa2, Hspa4, Hspa4l, Hspa5, Hspa8, Hsph1 | Mus musculus (Mouse) |
| Intersectin | E9Q0N0, B2RR82 | Itsn1, Itsn2 | Mus musculus (Mouse) |
| Kcp | Q3U492 | Kcp | Mus musculus (Mouse) |
| Lactate Dehydrogenase | P16125 | Ldhb | Mus musculus (Mouse) |
| Lrp1 | Q91ZX7 | Lrp1 | Mus musculus (Mouse) |
| Maged1 | Q9QYH6 | Maged1 | Mus musculus (Mouse) |
| Map7 | A2AG50 | Map7d2 | Mus musculus (Mouse) |
| Myosin | Q3UH59, Q69ZX3, E9Q174 | Myh10, Myh11, Myo6 | Mus musculus (Mouse) |
| Numb | Q9QZS3, O08919 | Numb, Numbl | Mus musculus (Mouse) |
| Osbp | D3YTT6 | Osbpl3 | Mus musculus (Mouse) |
| Pdcd | Q9WU78 | Pdcd6ip | Mus musculus (Mouse) |
| Phgdh | Q61753 | Phgdh | Mus musculus (Mouse) |
| Picalm | Q7M6Y3 | Picalm | Mus musculus (Mouse) |
| Pik3 | Q61194 | Pik3c2a | Mus musculus (Mouse) |
| Prune | Q52KR3 | Prune2 | Mus musculus (Mouse) |
| Periaxin | O55103 | Prx | Mus musculus (Mouse) |
| Reps | O54916, B9EI38 | Reps1, Reps2 | Mus musculus (Mouse) |
| Rundc3a | O08576 | Rundc3a | Mus musculus (Mouse) |
| Sec Faily | Q9D1M0, E9QAT4, Q01405 | Sec13, Sec16a, Sec23a | Mus musculus (Mouse) |
| Sh3d19 | Q91X43 | Sh3d19 | Mus musculus (Mouse) |
| Slc12a4 | Q9JIS8 | Slc12a4 | Mus musculus (Mouse) |
| Snap91 | Q61548 | Snap91 | Mus musculus (Mouse) |
| Alpha-synuclein | O55042 | Snca | Mus musculus (Mouse) |
| Sorting nexin | Q91VH2 | Snx9 | Mus musculus (Mouse) |
| Sos1 | Q62245 | Sos1 | Mus musculus (Mouse) |
| Spectrin Beta | Q62261 | Sptbn1 | Mus musculus (Mouse) |
| Stambpl1 | Q76N33 | Stambpl1 | Mus musculus (Mouse) |
| Ston2 | E9PXP7 | Ston2 | Mus musculus (Mouse) |
| Synj1 | D3Z656 | Synj1 | Mus musculus (Mouse) |
| Tfg | Q9Z1A1 | Tfg | Mus musculus (Mouse) |
| Tnrc6b | Q8BKI2 | Tnrc6b | Mus musculus (Mouse) |
| Tom1l2 | Q5SRX1 | Tom1l2 | Mus musculus (Mouse) |
| Trim21 | Q3U7K7 | Trim21 | Mus musculus (Mouse) |
| Tubulin | P68368, Q9CWF2, Q922F4 | Tuba4a, Tubb2b, Tubb6 | Mus musculus (Mouse) |
| Ubiquitin C | P0CG50 | Ubc | Mus musculus (Mouse) |
| Utrophin | E9Q6R7 | Utrn | Mus musculus (Mouse) |
| Vcp | Q01853 | Vcp | Mus musculus (Mouse) |

| | | | |
|---|---|---|---|
| Ybx | P62960, Q9JKB3 | Ybx1, Ybx3 | Mus musculus (Mouse) |
| Ywhag | P61982 | Ywhag | Mus musculus (Mouse) |
| FLJ45252 homolog | Q6PIU9 | FLJ45252 homolog | Mus musculus (Mouse) |
